# *In vivo* validation of spray-dried mesoporous bioactive glass microspheres acting as prolonged local release systems for BMP-2 to induce bone regeneration

**DOI:** 10.1101/2020.03.21.001404

**Authors:** Julia C. Berkmann, Aaron X. Herrera Martin, Carlotta Pontremoli, Kai Zheng, Christian H. Bucher, Agnes Ellinghaus, Aldo R. Boccaccini, Sonia Fiorilli, Chiara Vitale-Brovarone, Georg N. Duda, Katharina Schmidt-Bleek

## Abstract

Despite years of diligent research in fracture healing, an unmet clinical need for safe and effective pharmacological treatments to improve bone regeneration persists with 10 – 20 % of fracture cases exhibiting impaired healing. Bone morphogenetic protein-2 (BMP-2) is a known key mediator of physiological bone regeneration and is clinically approved for selected musculoskeletal interventions. Yet, broad usage of this growth factor is impeded due to side effects that are majorly evoked by high dosages and burst release kinetics.

In this study, mesoporous bioactive glass microspheres (MBGs) produced by an aerosol assisted spray-drying, scalable process were found to be biocompatible and to induce a pro-osteogenic response on human MSCs *in vitro*. Loading of the MBGs with BMP-2 resulted in prolonged, low-dose BMP-2 release without affecting the material features. In a pre-clinical rodent model, BMP-2 loaded MBGs significantly enhanced bone formation and influenced the microarchitecture of newly formed bone. The MBG carriers alone performed equal to the untreated (empty) control in most parameters tested, while additionally exerting mild pro-angiogenic effects.

Using MBGs as a biocompatible, pro-regenerative carrier for local and sustained low dose BMP-2 release could limit side effects, thus enabling a safer usage of BMP-2 as a potent pro-osteogenic growth factor.

## Introduction

Bone is among the few tissues in the adult mammalian organism that can fully regenerate and restore its physiological function (*restitutio ad integrum*) [1]. However, in the clinical routine, 10-20 % of fracture patients present with impaired bone healing, leading to delayed healing progression or non-unions [2, 3]. Not surprisingly, impaired bone healing strongly impacts the patients’ quality-of-life due to a prolonged time-to-recovery, additional surgical interventions and restricted physical activity. All factors oftentimes correspond to productivity losses as well as exacerbate the overall socio-economic burden [4].

The capacity for complete bone healing is known to decline with increasing age [5, 6], a consequence of a higher prevalence of comorbidities and a generally more pro-inflammatory state (“inflamm-aging”) [7]. The total number of impaired healing cases as well as its prevalence are expected to rise in light of the demographic change and a continuous increase in life expectancy [8-10], resulting in a growing medical need for strategies to restore healing in compromised settings.

Aside from autologous iliac crest bone grafting (ICBG), representing the gold standard treatment for severely impaired bone healing [11, 12], another prominent solution to restore healing is the pharmacological intervention with potent pro-regenerative drugs. Amongst these, proteins of the bone morphogenetic protein (BMP) [13] family, belonging to the TGF-β superfamily, have shown substantial potential to enable and accelerate bone regeneration. First reports on such effectiveness date back to 1889 [14]. Recombinant human BMP-2 (rhBMP-2) (INFUSE® Bone Graft, Medtronic Spinal and Biologics, TN; in Europe InductOS®, Medtronic BioPharma, NL) as well as BMP-7 (OP-1, Stryker Biotech, MI) received clinical approval for specific interventions, such as open fractures of the tibia or interbody spinal fusion [15], however, currently only rhBMP-2 products are available on the market [12]. Despite the narrow therapeutic applications BMP-2 is approved for, off-label uses have been frequently observed [16, 17]. At the same time, numerous reports of partly severe side effects, ranging from osteolysis and infections, immune reactions, potential carcinogenic effects as well as heterotopic/ectopic bone formation [12, 15, 17], have limited the routine clinical use [18, 19].

Clinically, BMP-2 is administered in conjunction with an absorbable bovine type-1 collagen sponge (ACS) onto which the BMP-2 solution is applied, leading to BMP-2 doses around 10-12 mg for treatments of long bone defects [20, 21]. Both, the ACS serving as a carrier as well as the supraphysiological dose of BMP-2, natively occurring in cortical bone in the range of 1-2 µg/kg [22], can be considered suboptimal [15].

Strikingly, the ACS itself has been found to significantly affect bone healing in a pre-clinical study, tested with the commercially available Helistat® (Xemax Surgical Products, Napa, CA) that is used for the administration of rhBMP-2. The ACS was found to cause strong immune responses as well as to impact the osteogenic potential and viability of human mesenchymal stromal cells (hMSCs) [23]. In the same study, another commercially available ACS (Lyostypt®, B. Braun, Germany) that yielded less severe *in vitro* immune and hMSC responses was observed to impair callus mineralization upon implantation in a 0.7 mm osteotomy gap in mice. Applying a collagen sponge in a pre-clinical model of impaired fracture healing by creating a critical-sized defect, the authors of the present study confirmed this finding (Supplemental figure S1). Moreover, it is known that collagen sponges possess rather unfavorable release kinetics, including low drug retention and high burst release [24, 25]. The poor release characteristics result in supraphysiological dosages that have to be applied to render BMP-2 available in sufficient amount at the implantation site despite rapid diffusion from implants and short half-life caused by proteolytic degradation [24, 25]. Thus, following the famous quote from Paracelsus “All things are poison, and nothing is without poison, the dosage alone makes it so a thing is not a poison” [26], it was postulated that the supraphysiological BMP-2 dosages are key contributors to the side effects [17]. Therefore, it has been suggested to realize strategies to apply minimal effective BMP-2 dosages [15].

Taken together, a reduction in dosage and/or an improvement of release kinetics can potentially dampen the side effects [25, 27] and lay the foundation for a broader and safer use of BMP-2. Accordingly, a carrier that features low burst and prolonged release, leading to lower effective dosages per time interval, as well as possesses inherent pro-regenerative properties, low immunogenicity and appropriate degradation kinetics would be an ideal candidate [18, 28].

In the current study, we analyzed mesoporous bioactive glasses, produced in the form of microspheres by aerosol assisted spray-drying method (SD-MBGs) [29], as an alternative promising carrier for BMP-2 instead of the currently used ACS. We selected SD-MBGs due to their intrinsic excellent bioactive behavior and related pro-regenerative potential, along with the high exposed surface area (approx. 200 m^2^/g) and regular nanopores (8-10 nm), which allow the storage and the release of active agents, such as drugs [30] or biomolecules [31]. In addition, the scalable and reproducible production route of SD-MBGs [29], and the possibility to impart multi-functionality by enriching the composition through the incorporation of selected therapeutic elements (e.g. strontium [32, 33]), enables a wide range of applications and can facilitate the clinical translation of the proposed carrier. Accordingly, we hypothesized that the SD-MBGs could act as a suitable carrier for BMP-2 in the bone healing context, exhibiting superior properties than clinically used carrier systems. We found that SD-MBG loading of rhBMP-2 did not alter the spherical morphology and the framework composition of SD-MBG. The release experiment of BMP-2 showed a low burst and a sustained release profile over the entire test interval. Overall, this release kinetics led to low dosages of solute growth factor in the range of 0.5 to 1 µg released BMP-2 over 14 days, which translates into 1-2 % of the clinically applied BMP-2 dosage (in humans 12 mg per application, based on comparison of dosage/kg). After validation of the carriers’ biocompatibility and pro-osteogenic effect on hMSCs and human blood, the carrier with and without BMP-2 load was embedded in an autologous blood clot, acting as a place-keeper for the MBG microspheres without delaying the progression of healing [18], and placed into a 2 mm osteotomy gap of a pre-clinical aged rat model of compromised healing [34]. Bone healing outcome was investigated radiologically, histologically and immunohistochemically, showing superior healing in the BMP group, while proving the suitability of the SD-MBG carriers alone to be utilized for bone regeneration purposes. With this approach we aim to provide evidence for the effectiveness and biocompatibility of bioactive glass-based BMP-2 carriers, potentially allowing more patients suffering from different fracture cases to benefit from the vast pro-osteogenic potential of this growth factor.

## Results

### 1. Ionic dissolution products of SD-MBGs are biocompatible on primary hMSCs

Before the evaluation of the spray-dried MBG microspheres with a binary SiO_2_-CaO composition [29] as drug carrier *in vivo* in an osteotomy model, the biocompatibility of SD-MBGs on primary hMSCs was studied *in vitro* in two concentrations (concentration 1 (c1) = 1.5 mg/mL, concentration 2 (c2) = 5 mg/mL) using a transwell assay. For this, the metabolic activity and cell number as well as lactate dehydrogenase (LDH) secretion of hMSCs from three donors in response to SD-MBG were considered (Figure 1). Since SD-MBGs were added into transwell inserts, the hMSCs were not in direct contact with the microspheres, but they experienced the change in ion concentration in the medium due to surface ion-exchange reactions and progressive dissolution of the MBG glass network. Both the relative metabolic activity (Figure 1A) and the relative cell count (Figure 1B) were found to be only slightly reduced for SD-MBGs (especially for c1) at day 3 and 5. In order to provide insights into the viability (here defined as metabolic activity) per cell, the cell vitality index was calculated by building a ratio of metabolic activity (Figure 1A) and cell count (Figure 1B). No decrease in cell vitality could be seen for cells treated with either concentration of SD-MBG at all time points tested (Figure 1C), which clearly indicates that the material is highly biocompatible. This finding is also supported by the results of the LDH cytotoxicity assay, since no significant increase in secreted LDH could be measured in any of the test groups compared to fresh culture medium (Figure 1D). Further evidence for this high biocompatibility was obtained by exposing human whole blood to SD-MBGs for 4 and 24 hours, which resulted in negligible secretion of the pro-inflammatory cytokines TNF-α, while IFN-γ could not be observed in the supernatants of SD-MBG treated whole blood (Supplementary figure S2).

**Figure 1.**
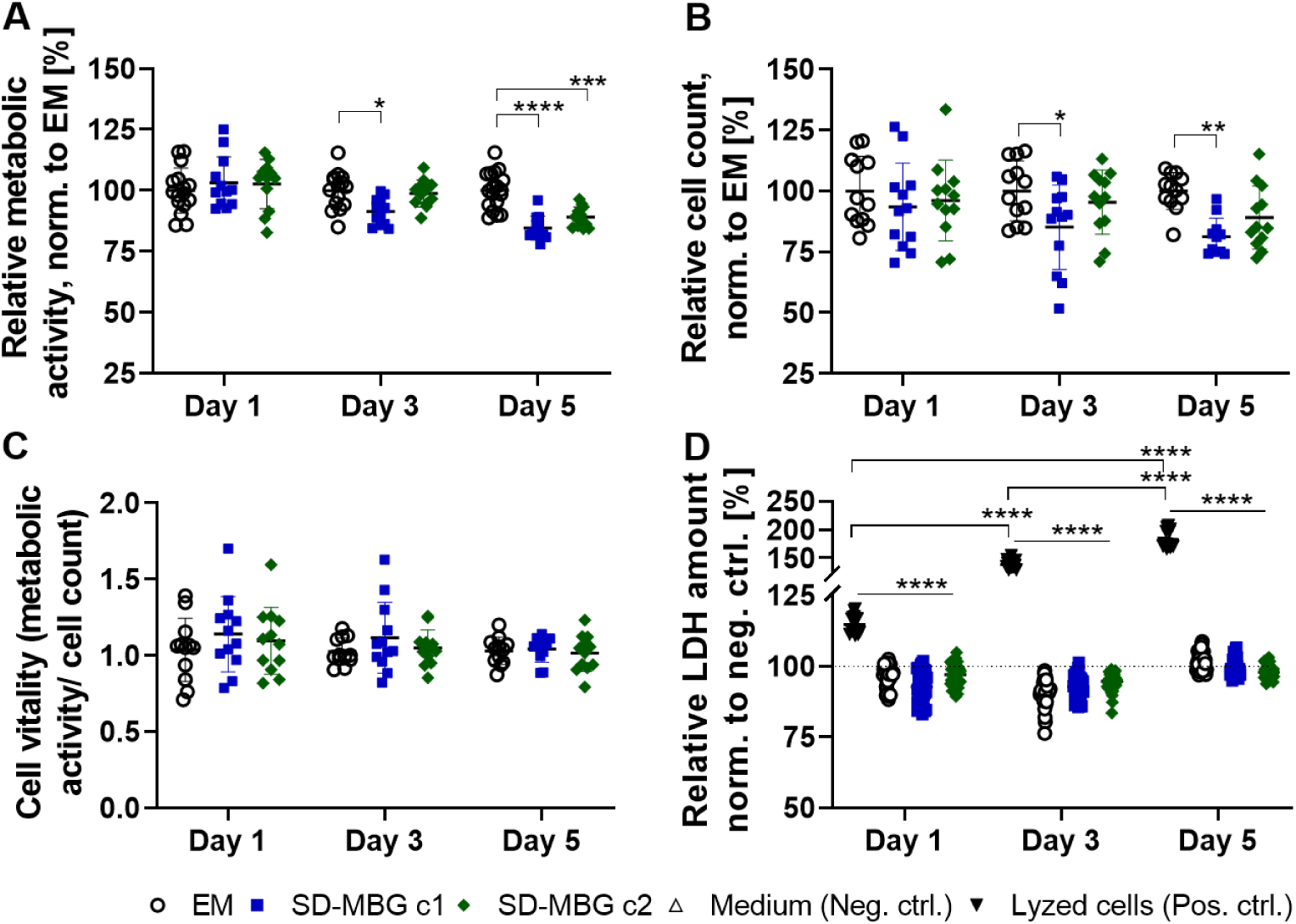
Biocompatibility and cytotoxicity upon indirect exposure of hMSCs to SD-MBGs for 1, 3 and 5 days. **A:** metabolic activity using Presto Blue and measuring substrate conversion by the cells. **B:** cell count as determined by DAPI staining, imaging of the wells (two 3×3 mosaic images per well using 10X magnification taken at slightly ex-centrical positions) and counting the nuclei using FIJI ImageJ software. **C:** cell vitality calculated by building a ratio of metabolic activity and cell count, indicating the viability per cell. **D:** Relative LDH content in the supernatant compared to fresh culture medium serving as negative control and set to 100 %. Cells cultured in EM were lyzed at each testing time point and were used as positive control to estimate the maximal amount of LDH that could be secreted into the supernatant. Normalization per time point as stated in the axis titles, n= 3 hMSC from different donors with 4 technical replicates each. ANOVA with Tukey’s multiple comparison test was performed. EM: cells cultured in expansion medium, SD-MBG: spray-dried mesoporous bioactive glasses, c1: concentration 1 (1.5 mg/mL), c2: concentration 2 (5 mg/mL).

### 2. hMSCs respond with higher osteogenic potential to SD-MBGs ionic extracts

After confirming the high biocompatibility of SD-MBGs on primary human MSCs, we aimed to validate the pro-osteogenic potential of the material on the same cell source. For this, osteogenesis was induced *in vitro* by the addition of pro-osteogenic additives to the medium. Similar to the biocompatibility tests, SD-MBGs were added to transwells and the hMSCs were exposed to ionic extracts from SD-MBGs over the entire course of the experiments. Using the same initial concentrations of SD-MBGs as in the biocompatibility experiments (1.5 and 5 mg/mL), matrix mineralization was studied by Alizarin Red staining after 10 and 14 days (Figure 2A) while collecting the supernatant before each medium change to quantify the amount of free phosphate released into the medium (Figure 2B). Matrix mineralization relative to the cell number was significantly increased when cells were exposed to the higher concentration (c2) of SD-MBGs, while the amount of free phosphate did not vary strongly across groups. hMSCs from three donors, previously tested and selected for their reduced osteogenic differentiation capacity (Supplementary figure S3), were exposed to the higher concentration of SD-MBGs. At day 14, hydroxyapatite deposits and cell nuclei were stained using OsteoImage and DAPI (Figure 2C). The intrinsically slow mineralization potential of the selected hMSCs was confirmed by the low amount of OsteoImage-positive areas (staining hydroxyapatite, green dots, marked with arrows). The mineralization-boosting effect of SD-MBGs in turn was evidenced by the substantially more pronounced OsteoImage signal. Altogether, these findings underline the pro-regenerative/ pro-osteogenic capacity of SD-MBG, making it an attractive candidate as a BMP-2 carrier for *in vivo* bone healing applications.

**Figure 2.**
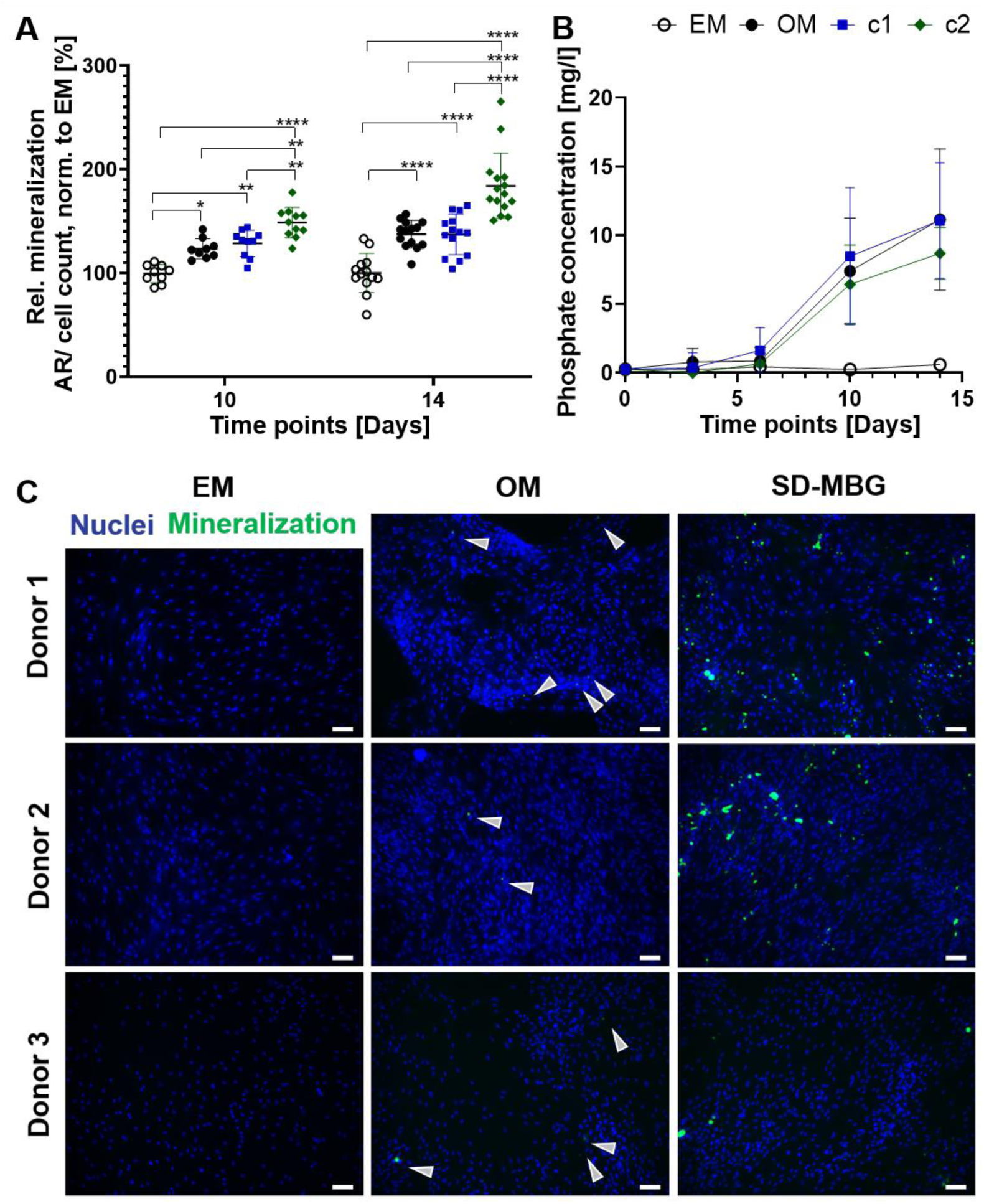
Matrix mineralization and free phosphate accumulation in supernatant of hMSCs upon osteogenic induction and continuous exposure to SD-MBG dissolution products over 14 days. **A:** Osteogenic differentiation induction in hMSCs by culture in osteogenic medium (OM) and exposure to SD-MBG ionic extracts. Relative matrix mineralization depicted as Alizarin Red (AR) optical density over cell count (DAPI signal), normalization to cells cultured in expansion medium (EM) (negative control). Significant matrix mineralization can be observed in all groups receiving OM (OM, concentration 1 (c1) = 1.5 mg/mL SD-MBGs and concentration 2 (c2) = 5 mg/mL SD-MBGs), with SD-MBG in c2 inducing significantly higher mineralization compared to all other groups. n= 3 hMSC donors with ≥ 3 technical replicates each. ANOVA with Tukey’s multiple comparison test was performed. **B:** Quantification of free phosphate accumulation in supernatant over time. In all groups receiving OM, an increase in phosphate amount can be observed starting at day 6. n= 3 hMSC donors with 3 technical replicates. **C:** At day 14 of osteogenic induction, OsteoImage staining (green) combined with nuclei staining (DAPI, blue) was conducted, imaged at 10x, scale bar = 100 µm. Representative images are shown for n= 3 hMSC donors. Arrows in the OM group point at mineralized nodules. Note the substantially increased matrix mineralization for the SD-MBG group, here shown as representative images from c2.

### 3. Characterization of SD-MBGs loaded with BMP-2

#### 3.1. Successful BMP-2 loading of SD-MBGs

SD-MBGs were successfully loaded with BMP-2 by incipient wetness impregnation method, as confirmed by differential thermal analysis (DTA) (Figure 3A). While no peaks were observed for the SD-MBG thermogram, SD-MBG loaded with BMP-2 (SD-MBG + BMP-2) showed a characteristic endothermic peak ascribed to the protein denaturation [35]. Additional proof for successful incorporation of BMP-2 was provided by thermogravimetric analysis (TGA) (Figure 3B). A negligible weight loss in the range of 25-150 °C ascribed to the release of the surface adsorbed water was observed for SD-MBG, validating the absence of any residual organic compounds. In comparison, a significantly higher weight loss in the same temperature range was detected for the SD-MBGs + BMP-2, due to the release of water bound to the BMP-2 protein. The additional weight loss in the temperature range between 200 and 400°C, observed exclusively for the BMP-2 loaded material, can be assigned to the decomposition of BMP-2. Moreover, zeta-potential analysis showed a negative surface charge (−24.5 ± 1.7) for SD-MBG particles alone when suspended in water, which resulted to be more negative after BMP-2 loading (−32.1 ± 1.9 for BMP-2).

**Figure 3.**
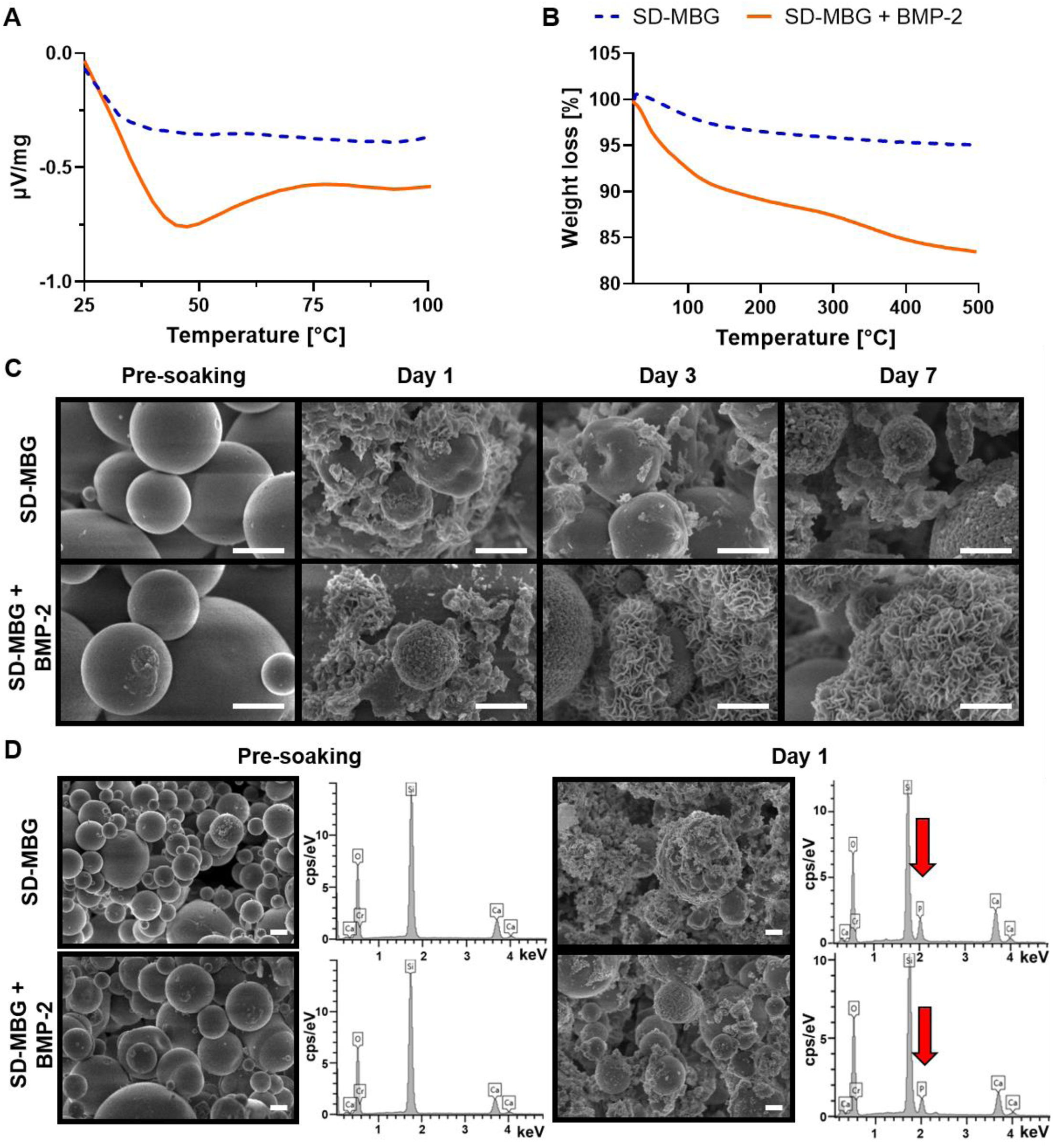
Thermoanalytical and morphological characterization of SD-MBG and SD-MBG + BMP-2. **A:** differential thermal analysis (DTA) and **B:** thermogravimetric analysis (TGA) showing an endothermic peak and pronounced weight loss only for SD-MBG + BMP-2, respectively, thereby confirming the loading of the protein. **C:** Field Emission Scanning Electron Microscopy (FE-SEM) of SD-MBG (top) and SD-MBG + BMP-2 (bottom) pre-soaking and at 1, 3 and 5 days of soaking in simulated body fluid (SBF). Starting at 1 day of soaking, hydroxyapatite formation can be detected by increasing roughness of the material surface and **D:** Energy-dispersive X-ray spectroscopy (EDS) analysis. Only after soaking in SBF, the EDS spectra yield a peak for phosphate (red arrows). Scale bar = 1 µm.

### 3.2. Morphological characterization of SD-MBG before and after BMP-2 loading

Concerning the morphology of SD-MBG and SD-MBG + BMP-2, Field Emission Scanning Electron Microscopy (FE-SEM) observations of SD-MBG and SD-MBG + BMP-2 (Figure 3C and D, dry particles pre-soaking in simulated body fluid (SBF)) showed spherical microspheres in the range of 1-5 μm. Nitrogen adsorption analysis revealed a specific surface area of 175 m^2^/g and an average pore size distribution around 8-10 nm (Supplementary figure S4), in fair agreement to the data previously published [29]. Energy-dispersive X-ray spectroscopy (EDS) spectra of both particles (Figure 3D, dry particles pre-soaking in SBF)) revealed a Si/Ca molar ratio in good agreement with the nominal one. The FE-SEM observations and EDS analysis of dry material powders (Figure 3C-D) evidenced that the BMP-2 loading into the mesopores does not alter the morphological features and the chemical composition of the MBG microparticles. In particular, the Si/Ca molar ratio revealed by EDS before and after the BMP-2 incorporation (Figure 3D) remained unaffected, indicating that the loading procedure did not induce substantial ion release.

#### 3.3. Bioactivity of SD-MBG alone and after BMP-2 loading

With the aim to investigate the bioactive behavior of SD-MBG alone and after loading with BMP-2, the microspheres were immersed in SBF and their ability to induce the formation of a hydroxyapatite layer on their surface was studied (Figure 3C and D). As highlighted by FE-SEM and EDS analysis, performed on dried powders collected after soaking, hydroxyapatite formation occurred after only 1 day of soaking, resulting in a compact layer of needle-like nanocrystals [36] covering the particle surface. EDS analysis (Figure 3D), revealed the presence of phosphorous and a Ca/P ratio very close to 1.7, the typical value reported in the literature for carbonated hydroxyapatite [37]. The agglomerates of the hydroxyapatite-like phase increased in size during the test, finally causing the full coverage of the surface (Figure 3C, day 7). These results clearly confirmed that the BMP-2 loading did not affect the SD-MBG bioactivity, allowing to preserve an essential feature of this material for application in bone regeneration processes [38]. Although it has been reported in the literature that bioactive glasses can induce a fast increase in pH when immersed in medium [39], for our samples, the pH of the SBF solution resulted in values below 7.8 during the entire *in vitro* bioactivity test, values that allow osteoblasts to maintain their physiological activity [40]. Taken together, BMP-2 was found to be successfully loaded into the mesopores of the SD-MBGs without altering the morphology, chemical composition or the intrinsic bioactivity of the carrier.

### 4. SD-MBGs enable prolonged BMP-2 release

The release kinetics of BMP-2 from the SD-MBG microspheres was investigated by soaking the BMP-2 loaded SD-MBG in either PBS or Tris-HCl with a physiological pH of 7.4 at 37 °C for 14 days. While the phosphate ions contained in PBS allow hydroxyapatite deposition, Tris-HCl medium prevents hydroxyapatite formation due to a lack of phosphates. Hence, an effect of hydroxyapatite formation on BMP-2 release kinetics can be clearly identified. The supernatant was collected repeatedly, serving as samples for BMP-2 quantification by means of enzyme-linked immunosorbent assay (ELISA) (Figure 4A). Overall, irrespective of the elution buffer utilized, a prolonged and sustained release of low amounts of BMP-2 was observed over the entire testing interval of 14 days, without an initial burst release. A cumulative BMP-2 release of 0.43 and 0.90 µg/mL per *in vivo* dosage (0.75 mg MBGs) of loaded SD-MBG was detected for elution in PBS and Tris-HCl, respectively.

**Figure 4.**
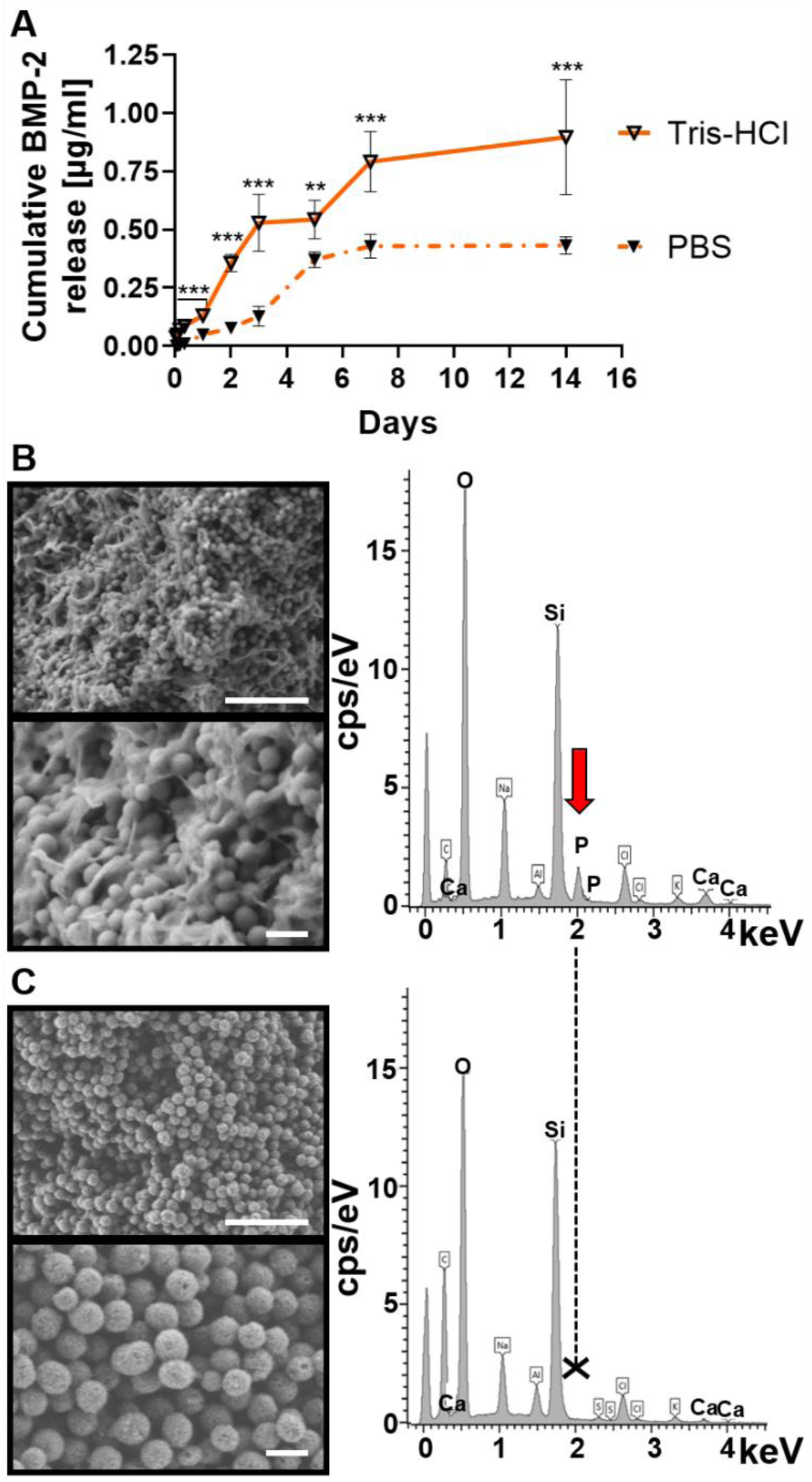
*In vitro* BMP-2 release experiment in PBS and Tris-HCl quantified via an α-BMP-2 ELISA and analyzed SD-MBG at the end of the release experiment by FE-SEM and EDS. **A:** BMP-2 cumulative release profile over 14 days, supernatants from SD-MBG + BMP-2, immersed in either PBS or Tris-HCl were sampled used for BMP-2 quantification. n=3 independent samples, tested at least in duplicates in the ELISA. Mann-Whitney U test was performed to compare the cumulative amount of detected BMP-2 between elution in PBS and Tris-HCl at each time point. **B + C:** FE-SEM images (left) and EDS spectra (right) of particles after the release experiment, **B:** in PBS, **C:** Tris-HCl. The red arrow points at the phosphate peak which is only present for the PBS-soaked SD-MBG. Top image of B and C scale bar = 1 µm, bottom image scale bar = 200 nm. ELISA: Enzyme-linked immunosorbent assay, FE-SEM: Field Emission Scanning Electron Microscopy, EDS: Energy-dispersive X-ray spectroscopy.

The clinically applied dosage of 12 mg BMP-2 in humans properly adjusted to the rat model based on the body weight yields a dosage of 50 µg BMP-2 per animal, already applied in previous studies [41, 42]. Accordingly, the amount of *in vitro* released BMP-2 over 14 days adds up to 1-2 % of the clinically used dosage. The approximately twofold higher release in Tris-HCl compared to PBS can be ascribed to the formation of hydroxyapatite deposits on the surface of the SD-MBG immersed in PBS, partially hindering the BMP-2 diffusion out of the pores (Figure 4B and C). While a phosphate peak appeared in the EDS spectra of SD-MBG soaked in PBS (Figure 4B, red arrow), indicative for hydroxyapatite formation, and the material surface exhibited the classical cauliflower morphology, both observations were nonexistent after soaking the particles in Tris-HCl (Figure 4C). The SEM images show a clear size reduction of the particles due to their gradual dissolution after immersion in both buffers if compared to the pre-soaking size of 1-5 µm (Figure 4B and C).

### 5. *In vivo* BMP-2 release from SD-MBG is beneficial for the bone healing process

#### 5.1. Radiological evaluation: BMP-2 release induces higher bone and tissue formation

The *in vivo* response induced by SD-MBG microspheres alone and in combination with BMP-2, showing a low dose release of approximately 0.5 – 1 µg BMP-2 over 14 days *in vitro* (Figure 4A), was evaluated in a femoral osteotomy model of compromised healing in female rats, as previously described by Preiniger *et al*. [34]. After creating a 2 mm bone defect in the femoral shaft, the animals received a hybrid formulation comprised of SD-MBG without or with BMP-2, finely dispersed in an autologous blood clot (BC). The BC was created by taking autologous blood, combined with SD-MBG depending on the test group, and coagulation of the blood occurred within a standardized mold to achieve comparable shapes of the BC. One group of animals did not receive any material (untreated/empty), another received the BC alone, both serving as a control groups. The healing progress was monitored radiologically by *in vivo* micro-computed x-ray tomography (µCT) analysis at two and four weeks post surgery. An increase in both, bone volume (BV) and tissue volume (TV) was observed over time for all groups, with the SD-MBG + BMP-2 group exhibiting the highest values (Supplementary figure S5), already indicating *in vivo* the beneficial effect of the prolonged, low-dose BMP-2 treatment regime. After finalization at four weeks, the osteotomized bones were prepared for analysis by *ex vivo* µCT, histology and immunohistochemistry. The reconstructed µCT images were used to study the bridging stage of the fracture gap by blinded image evaluation. For this, µCT images from two planes of all animals (Supplementary figure S6) were presented to four evaluators in a blinded fashion, asking them to categorize the fracture gaps showing “no bridging”, “partial bridging” or “complete bridging”, a scoring previously described [43]. Within the BMP-2 group, complete bridging occurred in 50% of cases, in the other 50%, at least partial bridging was observed. For the vast majority of all other animals, no bridging was seen (Table 1).

**Table 1.** Results of the evaluation of bridging states. 1= complete bridging, 2= partial bridging, 3= no bridging. Only for the BMP-2 group, partial or complete bridging was observed in all animals. N= 4-6 animals per group. Scoring values stated as median form four different evaluators and summarized as the median of all animals per group. Empty: untreated control, BC: blood clot, SD-MBG: spray-dried mesoporous bioactive glass; BMP-2: bone morphogenetic protein 2.

|  | Empty | BC | BC + SD-MBG | BC + SD-MBG + BMP-2 |
| --- | --- | --- | --- | --- |
| **Animal 1** | 3 | 3 | 3 | 1 |
| **Animal 2** | 3 | 3 | 3 | 2 |
| **Animal 3** | 2,5 | 3 | 3 | 2 |
| **Animal 4** | 3 | 3 | 3 | 1 |
| **Animal 5** | 3 | x | x | x |
| **Animal 6** | 2 | x | x | x |
| **Median all animals** | **3,0** | **3,0** | **3,0** | **1,5** |
| **Bridging** | **No** | **No** | **No** | **Complete/partial** |

The analysis of *ex vivo* µCT data (Figure 5), which allows higher resolution imaging than the *in vivo* µCT, confirmed the significantly higher bone and tissue formation for the SD-MBG + BMP-2 compared to all other groups (Figure 5A). The ratio of bone volume within the callus volume (BV/TV) was not significantly affected by the BMP-2 treatment, since both BV and TV were elevated in this group. Bone mineral density (BMD) for the BMP-2 group was found to be decreased (Figure 5A) in the volume of interest (VOI), which also includes cortical bone fragments. This can be explained by the larger callus volume that was formed in the BMP-2 group, resulting in more newly formed bone when compared to the other groups with less callus volume, where the cortical bone represents a higher percentage of the total volume. However, upon excluding the cortical bone fragments, no significant change in callus BMD could be observed between groups (*in vivo* µCT data, Supplementary figure S5C). Despite similar ratios of bone volume within the callus volume (BV/TV) in all groups, the mineral content within the callus volume, also called bone mineral content (BMC), is significantly increased upon BMP-2 treatment, indicating that a higher net amount of deposited hydroxyapatite can be found in the formed callus (Figure 5A). Comparing the SD-MBG groups with and without BMP-2 in respect to their microarchitecture, BMP-2 release into the fracture area decreased the trabecular thickness (Tb.Th.) and increased the trabecular number (Tb.N.), thus inducing a callus with a finer microarchitecture, but increased trabecular branching (Figure 5B). The polar minimal moment of inertia (MMI (polar)), a 3D computational calculation of torsional stability, showed a significantly higher value for the BMP-2 group, indicated a higher torsional stiffness as a consequence of the progressed healing under BMP-2 influence (Figure 5C). Examples for reconstructed µCT images are depicted in Figure 5D.

**Figure 5.**
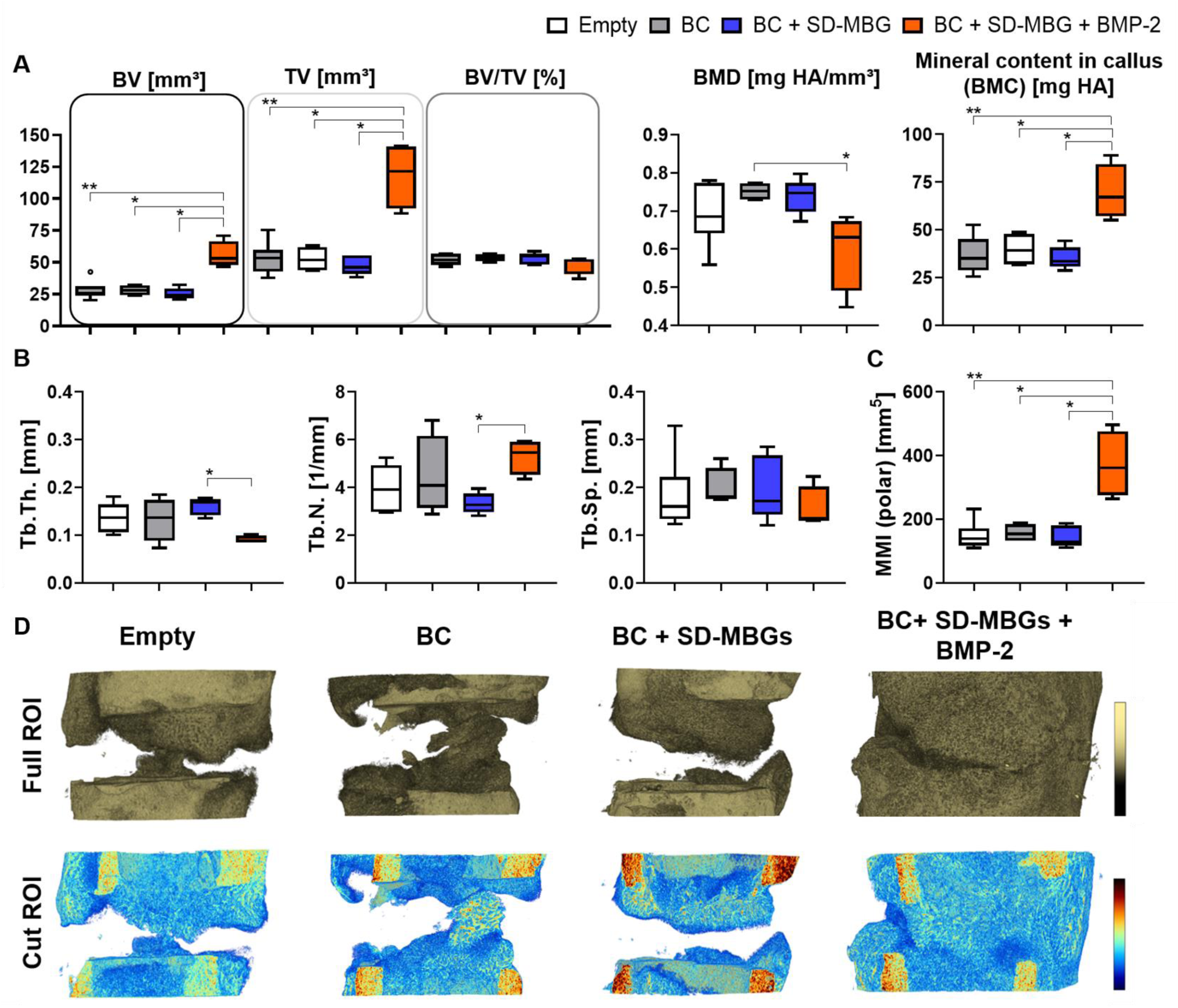
Ex vivo µCT analyses of the rat femora osteotomy gap area at 4 weeks post surgery. **A:** Bone volume (BV) and tissue volume (TV) were increased in the BMP-2 group while no difference was observed across all other groups. The ratio of BV and TV (BV/TV) remained unaffected. The bone mineral density (BMD) was reduced for the BMP-2 group, while the bone mineral content (BMC, BMC= TV* BMD) within the callus increased. **B:** microarchitecture of the bone in the osteotomy area was similar among all groups, except for the BMP-2 group, exhibiting a lower trabecular thickness (Tb.Th.) and higher trabecular number (Tb.N.) compared to the SD-MBG group, while trabecular separation (Tb.Sp.) was not changed. **C:** Polar minimal moment of inertia (MMI (polar)) was significantly higher in the BMP-2 group, depicting a more stable callus to torsional load, compared to all other groups. **D:** representative images of 3D rendered images of the osteotomy area. Top panel showing the entire volume of interest (VOI), bottom panel depicting density distributions of a longitudinally sectioned half of the VOI. The color scale indicates the mineralization state which increases from blue to red. **A-C:** N= 4-6 animals per group, shown are Tukey box plot distributions with line at median. Mann-Whitney U test was applied using the BMP-2 group as comparator.

Based on the *in vivo* and *ex vivo* µCT results, an effective *in vivo* release of BMP-2 can be postulated, as in this group the bone healing outcome was significantly enhanced compared to the SD-MBG carrier alone. Upon comparing the BC alone and in combination with SD-MBGs, no radiological difference could be determined (Figure 5). Similarly, the comparison of either the BC alone or BC + SD-MBG with the empty group, did not yield significant differences. Thus, it can be assumed that neither the BC nor the addition of SD-MBG impair the healing outcome, rendering the tested hybrid system a suitable drug release platform.

#### 5.2. Histological and immunohistochemical evaluation: BMP-2 release advances healing stage

The histomorphometrical anaylsis (Figure 6A) of the harvested bones based on the MOVAT’s pentachrome staining (Figure 6B) revealed the highest cartilage amounts in the empty group with no differences between BC + SD-MBG and BC + SD-MBG + BMP-2. In line with the µCT analysis, the mineralized area per total area was significantly higher in the BMP-2 group, while connective tissue was mostly absent from the fracture gap in this group (Figure 6A). Moreover, Sirius Red staining was conducted, which stains collagen fibers and allows to distinguish the different collagen types under polarized light [44]. The Sirius Red staining clearly showed a more advanced healing stage for the BMP-2 treated group, presenting high amounts of collagen type 1 fibers that can be correlated to the woven bone having formed in the fracture gap as seen in the MOVAT’s pentachrome staining in yellow/orange (Figure 6B, also compare the magnified images, bottom panel).

**Figure 6.**
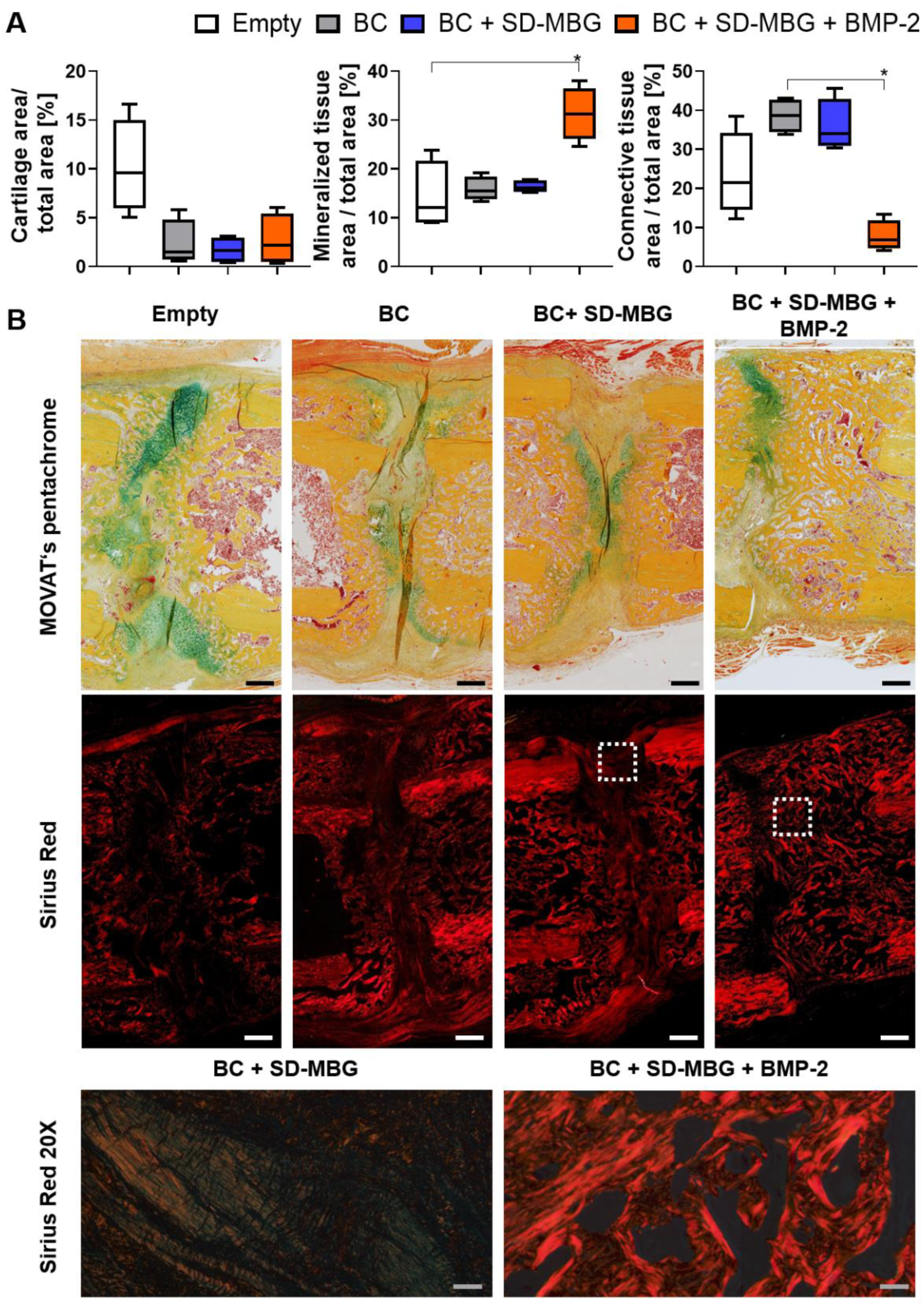
Histological and histomorphometrical analysis of rat femora at 4 weeks post osteotomy of the osteotomy gap area via MOVAT’s pentachrome and Sirius Red staining. **A:** Histomorphometry evaluation of the tissues present in the osteotomy gap based on the MOVAT’s pentachrome. The cartilage, mineralized and connective tissue fraction are depicted normalized to the callus area. Shown are Tukey box plot distributions with line at median. Mann-Whitney U test was performed using the BMP-2 group as comparator. **B:** Representative images of the gap area for all groups. Top panel: MOVAT’s pentachrome (yellow/orange: mineralized tissue, green/blue: cartilaginous tissue, orange/red: muscle tissue), 10X magnification. Middle and bottom panel: Sirius Red staining, imaged using polarized light. Middle panel: 10X magnification. Bottom panel: 20X magnification, imaged at the periosteal side in between the cortical bone fracture ends. Red stained fibers represent collagen type 1, greenish fibers represent collagen type 3. Scale bars for 10X = 500 µm, 20X = 50 µm.

Immunohistological analyses were conducted to quantify vessel formation and monocyte/ macrophage/ osteoclast cells in the callus area. α-SMA (α-smooth muscle actin) is a major component of the actin cytoskeleton of vascular smooth muscle cells, allowing contraction of the vessels [45] and this marker can be utilized to detect vessels in the osteotomy gap. Figure 7A depicts the quantification of α-SMA+ area over the standardized region of interest, together with the representative image of the callus and a magnified image of α-SMA+ areas (arrows). While no significant differences were found, the SD-MBG component in the BC + SD-MBG hybrid system appears to induce a higher α-SMA+ area in the fracture gap, indicating a higher pro-angiogenic effect and hinting at the *in vivo* pro-regenerative potential of the carrier alone. Immunohistological staining for CD68, a marker of the monocyte/ macrophage/ osteoclast lineage, was conducted to visualize bone turnover at the surface of the bone. We focused on CD68+ cells residing on the bone surface, which is an indicator for bone remodeling by osteoclastic cells (Figure 7B). The percentage of CD68+ cells residing on newly formed bone was quantified by measuring the surface (µm) of CD68+ cells that cover the mineralized callus fraction and the perimeter of the mineralized callus. No significant difference in fraction of mineralized bone covered by bone residing CD68+ cells could be determined across the groups, indicating that bone remodeling processes occur in all osteotomized bones irrespective of the treatment.

**Figure 7.**
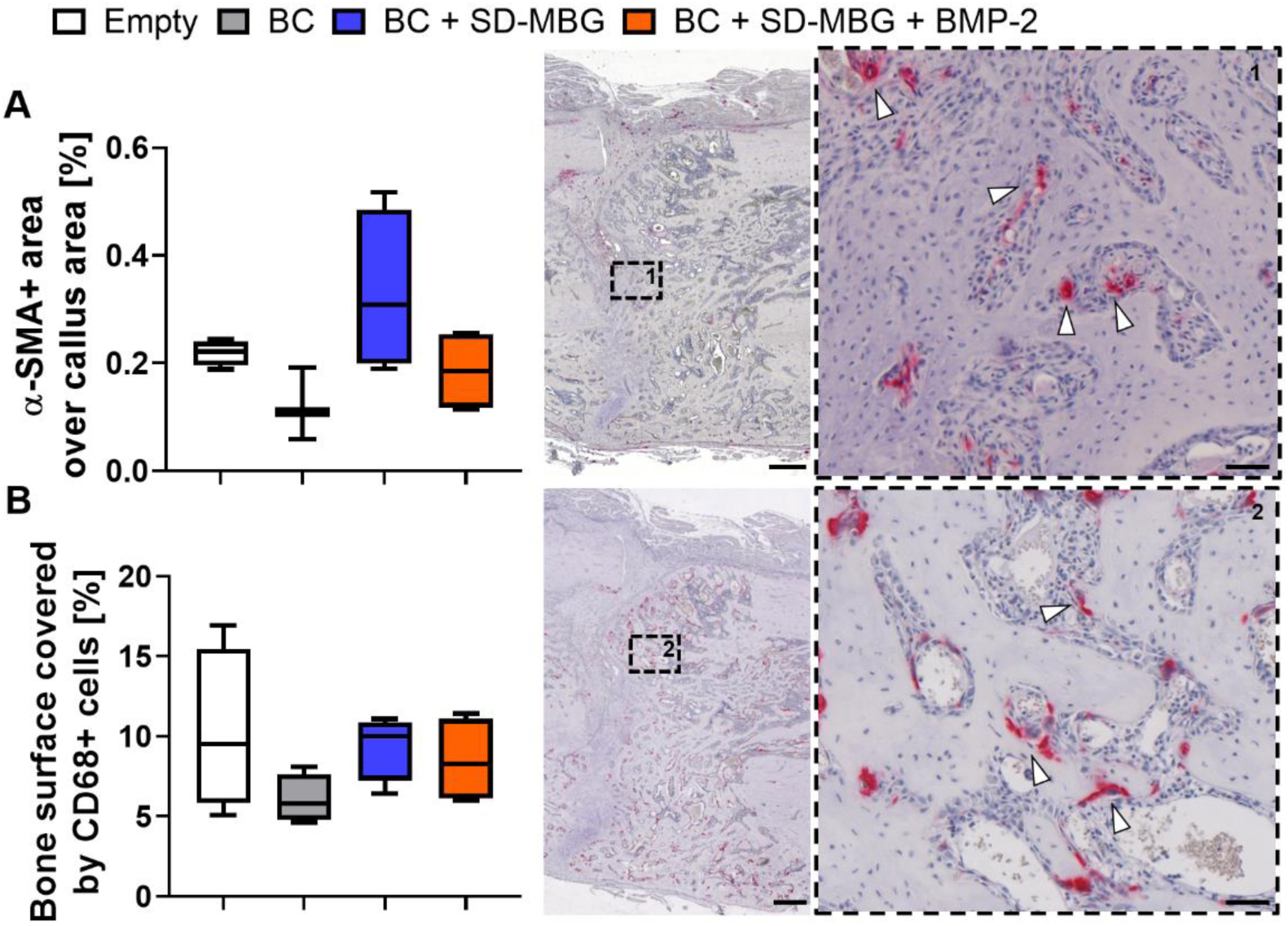
Immunohistochemical (IHC) analysis of rat femora at 4 weeks post osteotomy in the osteotomy gap area for α-SMA (A) and CD68 (B). **A, B** (left): Shown are Tukey box plot distributions with line at median; (middle and right panel): exemplary image of the region of interest (overview, 10x) as well as magnified image (40x, dashed box) showing the localization of the IHC positive cells for a selected region in the middle of the fracture gap are depicted. Hematoxylin counterstain was used for better tissue visualization (purple/blue). White arrows point at examples of the positive α-SMA+ and CD68+ staining (red). **A:** IHC staining of α-SMA for blood vessel quantification based on α-SMA+ area normalized to callus area. **B:** IHC staining of CD68 for quantification of osteoclastic cells based on the length of CD68+ cell contact to newly formed bone over total perimeter of mineralized callus. N= 4, scale bar= 500 µm in overview image (middle), scale bar= 50 µm in magnified image (right).

## Discussion

In this study, we validated spray-dried mesoporous bioactive glass microspheres (SD-MBG) [29], as a suitable carrier for prolonged, low-dose BMP-2 release, exerting beneficial effects on the bone healing outcome. In contrast to the conventional ACS as BMP carrier [23], we report for SD-MBG an excellent biocompatibility without a negative impact on bone formation when placed in the bone osteotomy gap. Moreover, cell vitality was not affected by SD-MBG when tested on primary human MSCs and the pro-inflammatory response from human whole blood was negligible. Ionic dissolution products of SD-MBG were revealed to even amplify the osteogenic differentiation of hMSCs *in vitro*, indicating the potential of being pro-regenerative in the context of bone healing. Loading SD-MBG with BMP-2 did not induce any effect on the material morphology and composition. SD-MBG showed no initial burst release while retaining a sustained low-dose release in the range of 1-2 % of clinically applied BMP-2 over the entire testing interval of 14 days (approximately 0.5 – 1 µg compared to 50 µg *in vivo* dosage in a rat osteotomy model [41, 42]). When applied in the fracture gap of a pre-clinical animal model of compromised healing [34], pure SD-MBG did not impair the healing progress, but rather exerted mild pro-angiogenic effects. The additional BMP-2 load was found to improve the healing outcome in all tested bone healing parameters, indicating that SD-MBG microspheres represent a suitable carrier and BMP-2 release platform for impaired bone healing scenarios, with the carrier possessing an intrinsic pro-regenerative potential (Table 2).

**Table 2.**
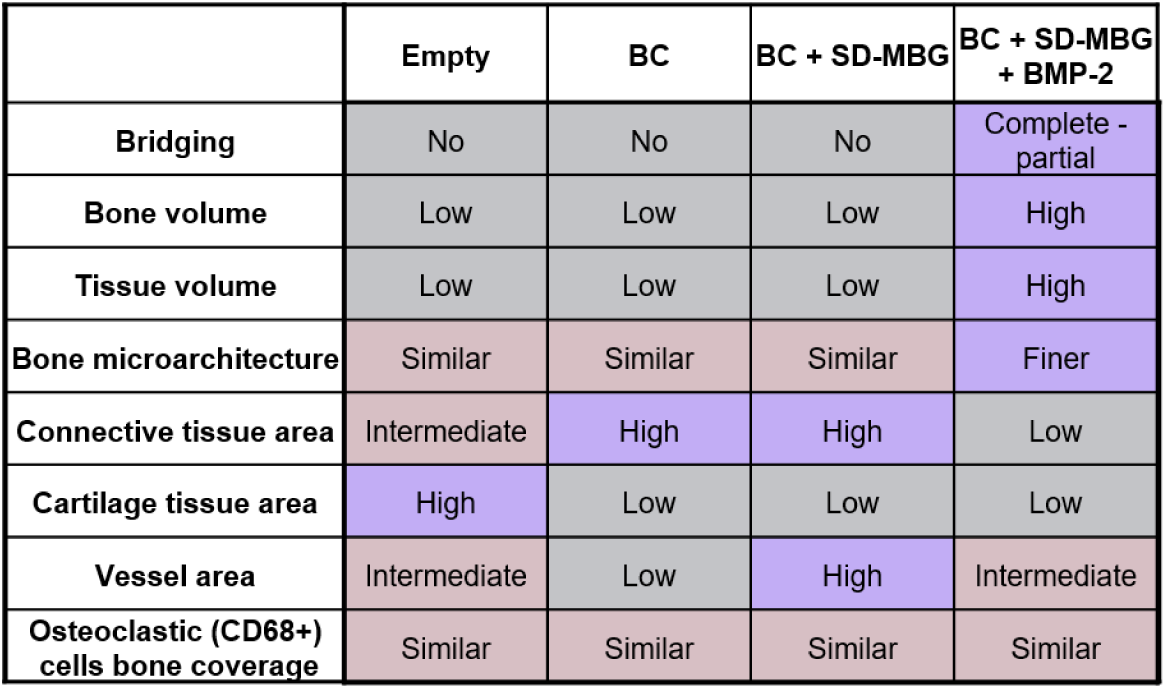
Tabular summary of the *in vivo* bone healing analysis. The color coding is based on the values obtained in the different analysis. Grey: absent bridging/ low values, reddish: intermediate values/ similar outcomes between groups, purple: observed bridging / finer callus and high values. Empty: untreated control, BC: blood clot, SD-MBG: spray-dried mesoporous bioactive glass; BMP-2: bone morphogenetic protein 2.

Initial clinical studies investigating the effects of BMP-2 administered via an ACS, including the large, multi-centered human “BMP-2 evaluation in surgery for tibial trauma” study (BESTT), showed superior effects of the treatment due to decreased rates of revision surgeries and infections with a 1.5 mg/mL BMP-2 dosage [20]. Other studies confirmed the beneficial BMP-2 effects while extending the range of application to other long bone defects [46]. As a result of these early studies, BMP-2 was increasingly applied in the clinics. Subsequently, reports on side effects largely neglected in the initial studies appeared, leading to a reassessment of the treatment’s safety and efficacy. Carragee *et al*. estimated that the treatment risk is 10 to 50 fold higher than originally anticipated [47]. There is vast agreement in the scientific and clinical community that most of the treatment-related risks arise from the supraphysiological dosage that needs to be applied [17], among others because of the unfavorable release kinetics of the clinically used collagen sponge [24, 25] with a burst release of ∼50 % within the first day post implantation [48].

Accordingly, other biomaterials acting as BMP-2 carrier have been investigated, ranging from inorganic materials to polymers (both synthetic and natural) to composites of such materials in various forms of delivery, as concisely summarized in the review by El Bialy *et al*. [12]. Indeed, it could be demonstrated that release systems exhibiting spatiotemporal controlled BMP-2 release kinetics evoked superior bone regeneration compared to the clinically used ACS [49] and allowed for lowering the dose significantly below the effective concentration if administered via the ACS [50]. Favorable release kinetics can reduce potential side effects as observed by Kowalczewski *et al*. by a decreased ectopic bone formation, while BMD at the mandibular defect site was increased upon comparing kerateine hydrogels with ACS [25]. Although a dose-response relationship has been reported for BMP-2 in the clinics with higher doses being more beneficial for the healing outcome [20], thorough testing of the optimal BMP-2 dosage regime unraveled that therapeutic concentrations can induce robust bridging with normal microarchitecture, while supraphysiological doses resulted in bridging as well, albeit causing lower bone quality with abnormal cyst-like bone structure [27]. These examples underscore that BMP-2 dosing and release kinetics are key factors for successful and safe therapy; suitable release kinetics can allow either the utilization of standard doses, or even a reduction in required BMP-2 amounts. At the same time, the required release kinetics might vary according to the pathology that needs to be treated and it is profoundly dependent on the carrier, the process of BMP-2 incorporation, interactions between protein and carrier and it’s physical configuration [51]. Focusing on the proper choice of biomaterials acting as BMP-2 carriers, the main requirements that an ideal biomaterial applied in the bone healing context should meet are biocompatibility, controlled degradation, an intrinsic pro-regenerative potential as demonstrated by either induction of vascularization or osteoconductivity/ osteoinductivity [18], a low immunogenicity and a suitable, controlled drug release profile [15]. Additionally, it is of outmost importance that a carrier exhibits high retention capacities and localizes the drug at the target site [12].

Bearing these criteria in mind, we employed SD-MBG with SiO_2_-CaO binary composition, mainly due to its excellent biocompatibility and sustained degradation upon exposure to physiological fluids [38, 52]. Enhancement of the pro-regenerative potential due to ionic dissolution products from bioactive glasses has been demonstrated on various cell types [38, 39, 53] and is highly dependent on the ionic composition, size and interaction mode [39, 54]. In this study, we were able to assess the SD-MBG intrinsic pro-osteogenic and osteoinductive properties *in vitro* and found indications for a pro-angiogenic effect *in vivo*. The pro-osteogenic effects mediated by SD-MBG could be explained by the release of Ca^2+^ ions that have been found to upregulate proteins belonging to the mitogen-activated protein kinase (MAPK) signaling pathway [55]. MAPK signaling in turn causes the phosphorylation of runt-related transcription factor 2 (RUNX2), a master transcription factor that mediates osteogenic lineage determination [56, 57]. The trend observed *in vivo* of increased vessel number in the blood clot group treated with embedded SD-MBG compared to animals only receiving the blood clot can likely be linked to the release of silicate species, which are reported in the literature to exert pro-angiogenic properties [58]. *In vitro* and *in vivo*, a low immunogenicity of the material was detected as well, distinguishing this material from other biomaterials that exert pro-inflammatory effects for example during their degradation as described for instance for poly(d,l-lactic-co-glycolic Acid, PLGA) [59, 60]. In the current study, a low-dose, sustained release profile of BMP-2 without burst effect from SD-MBG mesoporous structure, was found to effectively induce superior bone healing compared to the carrier alone. *In vitro*, a maximum cumulative release over 14 days of approximately 1 µg/mL was measured. In another rat study investigating bone healing, 1 µg was observed to be ineffective in promoting bone healing if administered via a collagen sponge [25], again highlighting the importance of proper release kinetics. The obtained release profiles (Figure 4A) depend on multiple factors, such as the occurrence of multiple interactions between proteins and internal pore surface, hydroxyapatite deposition partially blocking SD-MBG mesopores, as well as the overall morphological features. In particular, BMP-2 loading was driven by adsorption on MBG surface through the engagement of intermolecular interactions (e.g. mainly H-bonding) between surface –OH species and BMP-2 protein functionalities (−NH_2_, -COOH) as was described for BMP-2 binding to hydroxyapatite [61]. Furthermore, SD-MBG possess a negative surface charge, due to deprotonated silanols, able to bind positively charged protein molecules through electrostatic interactions. This charge-dependent binding was reported for BMP-2 binding the kerateine [62] or alginate carriers [63].

The degradation behavior of the carrier is highly relevant for the release properties. Albeit a lack of understanding of *in vivo* dissolution kinetics for the SD-MBG, we have clear evidence for their *in vitro* resorption. The particle size ranged from 1-5 µm pre-soaking, irrespective of the immersion medium, while after 14 days, the average size of the spheres was found to be below 200 nm, translating to a 5-to 25-fold size reduction (Figure 3, 4). In addition, the presence of phosphate ions in the surrounding solution causes hydroxyapatite formation on the surface of the SD-MBG (Figure 3, 4) which can partially block the mesopores leading to a dampened BMP-2 release compared to medium lacking phosphate ions (Figure 4A). It is noteworthy that the local *in vivo* concentration of phosphate ions is estimated to be 3-to 10-fold lower compared to the concentration found in PBS, this estimation is based on the determination of phosphate levels in human blood serum [39] and the evaluation of ion species abundance in the fracture hematoma analyzed in sheep (Supplementary figure S7). Lastly, the size of the spheres is essential in determining the degradation kinetics [39] since particle and pore size define the total exposed surface and thus the loading capacity.

The SD-MBG were embedded in an autologous blood clot for *in vivo* application and validation. This hybrid formulation can be easily produced even in the clinical context. In a previous study, the importance of the fracture hematoma for a successful fracture healing has been described [64], the autologous blood clot can be considered as an artificial but similar tissue when compared to the initial fracture hematoma. Both tissues are derived from blood, therefore contain similar cellular species, and underwent the process of coagulation. Other biomaterial-based approaches oftentimes spatially limit the formation of a hematoma due to hindrance/blockage of new tissue formation in the fracture area. In this approach however, the hematoma-like BC becomes an integral part of the composite. The administration of the BMP-2 loaded SD-MBG in particulate form enables versatility in dosing, since different clinical application certainly require adjustable concentrations, and shaping of the blood clot is equally customizable. The SD-MBG can be injected if contained in a liquid dispersion and the embedding moiety can be exchanged based on the specific requirements, e.g. to a thermo-sensitive hydrogel that solidifies at body temperature [30, 65]. However, it should be noted that the composite utilized in this study does not provide mechanical or distinct structural support to guide bone formation. Similar to the clinically used ACS, an additional support structure would need to be applied to ensure proper fixation of the fracture. In previous studies, we have demonstrated that the pro-regenerative potential of the MBG carriers depends on the composition [39] and can be steered based on the introduction of therapeutic ions into the glass network [32, 38]. Ion-doping opens several opportunities for synergistic drug-ion actions, thereby enabling potential multi-functionality. SD-MBG could be further enhanced by loading separately two or more drugs and combining them during their application. In this context, an *in vitro* study suggested that the combined use of BMP-2, −7 and −9 could significantly reduce the rate of BMP non-responders from 25-30 % to around 6 %. Lack of responsiveness to BMP treatment is a reported clinical challenge affecting up to 36 % of BMP-treated patients [19], thus a BMP combination therapy could prove beneficial to increase treatment efficacy. Moreover, the scalable production route of SD-MBGs, avoiding the use of hazardous and expensive solvents as well as inflammable solvents (e.g. ethanol) [29], can facilitate clinical translation of the proposed carrier.

Open questions remain concerning the optimal dosage and formulation of BMP-2 [66], potential synergistic effects using additional clinically approved drugs or therapeutic ions, and the most effective combination of loaded SD-MBG with mechanical support structures to ensure proper bone fixation.

## Conclusion

We developed and validated mesoporous bioactive glass (SiO_2_-CaO binary composition) in the form of microspheres as suitable BMP-2 carrier for enhancing/ restoring impaired bone healing cases. The favorable prolonged release of low-dose BMP-2 (around 1-2 % of the clinically applied dosage released over 14 days) was able to induce superior progression of the bone regeneration cascade. The distinct production route applying the aerosol spray-drying assisted method under mild aqueous conditions represents a scalable, cost and safety effective approach [29]. The combination of SD-MBG with an autologous blood clot for *in vivo* application has proved successful and could easily be translated to the clinical routine. These findings revealed a translatable biomaterial-based approach to limit side effects of BMP-2 usage by dampening excessive amount of soluble BMP-2 as observed for clinically employed collagen sponges, with the MBG possessing intrinsic characteristics that are beneficial for bone healing.

## Experimental Section

### Materials for the production of MBG samples

Pluronic P-123 (EO_20_PO_70_EO_20_, Mn ∼5800 Da), double distilled water (ddH_2_O), tetraethyl orthosilicate (TEOS) and calcium nitrate tetrahydrate (Ca(NO_3_)_2_ · 4H_2_O, 99 %), were purchased from Sigma Aldrich (St. Louis, USA) and used as received. All solvents were purchased from Sigma Aldrich (St. Louis, USA) in analytical grade.

### Synthesis of MBG Samples by Aerosol-Assisted Spray Drying Method

Based on the procedure reported by Pontiroli et al. [29], MBG micro-particles with a binary SiO_2_-CaO composition (molar ratio Si/Ca= 85/15 hereafter named SD-MBG) were synthesized by aerosol-assisted spray drying (SD) method. Briefly, 2.03 g of the non-ionic block copolymer Pluronic P123 were dissolved in 85.0 g of double distilled H_2_O (ddH_2_O). In a separate batch, 10.73 g of TEOS were pre-hydrolyzed under acidic conditions using 5.0 g of an aqueous HCl solution at pH= 2 until a transparent solution was obtained. The TEOS solution was then dropped into the Pluronic P-123 solution and kept stirring for 30 minutes. Thereafter, 1.86 g of calcium nitrate tetrahydrate were added. The final solution was stirred for 15 min and finally sprayed with a Mini Spray-Dryer B-290 (Büchi Labortechnik, Flawil, Switzerland) using nitrogen as the atomizing gas (inlet temperature 220 °C, N_2_ pressure 60 mmHg, feed rate 5 mL/min). The obtained powder was calcined at 600 °C in air for 5 h at a heating rate of 1 °C/min using a furnace (Carbolite 1300 CWF 15/5, Carbolite, Hope Valley, UK), in order to remove the templating agent.

### BMP-2 loading

SD-MBG were sterilized by heating at 160 °C for 2 h before BMP-2 immobilization. 500 μg BMP-2 (rhBMP-2, Peprotech, Hamburg, Germany) was dissolved in 1 mL of ultrapure water to form BMP-2 solution (500 μg/mL). 20 μL of BMP-2 solution was added to 0.75 mg of SD-MBG. After adsorption for 30 min, another 20 μL of BMP-2 solution was added. This process was repeated until 50 μg of BMP-2 were added to 0.75 mg of MBG particles, corresponding to the clinically applied total dosage of 12 mg BMP-2 in humans [20, 21]. For this, an average human body weight of 75 kg was assumed, leading to an average dose of BMP-2 of 0.16 mg/kg bodyweight. The body weight of the rats in this study was around 300 g, hence, 50 µg BMP-2 represent a fair approximation to the human dose. For *in vitro* analysis of bioactivity and morphology, 15 µg BMP-2/0.75 mg SD-MBG were loaded. The BMP-2 loaded SD-MBG (SD-MBG + BMP-2) were then dried overnight at 37 °C under sterile conditions.

### Characterization

#### FE-SEM and EDS

The morphology of SD-MBG and BMP-2 loaded SD-MBG (SD-MBG + BMP-2, 20 µg BMP-2 per mg SD-MBGs) particles was analyzed by Field-Emission Scanning Electron Microscopy (FE-SEM) using a ZEISS MERLIN (Carl Zeiss, Oberkochen, Germany) instrument. For FE-SEM observations, 5 mg of both SD-MBG and SD-MBG + BMP-2 powders were dispersed on a conductive carbon tape and coated with a chromium layer. Compositional analysis of the powders was performed by energy dispersive spectroscopy (EDS) using an AZtec EDS (Oxford instruments, Abingdon, United Kingdom). The spectra were collected on powders dispersed on carbon tape by analyzing an area of 75×50 µm.

#### Thermogravimetric and differential thermal analysis and Zeta-potential

The successful loading of the protein was assessed by thermogravimetric and differential thermal analysis (TG/DTA) and Zeta-potential measurements. TG/DTA were conducted on a STA 2500 Regulus (Netzsch-Gerätebau, Selb, Germany), collecting the thermograms over a temperature range of 25–500 °C with a heating rate of 10 °C/min under nitrogen in a flow of 40 mL/min. The Zeta-potential of SD-MBG and SD-MBG + BMP-2 in ultrapure water was measured using a Zetasizer Nano ZS (Malvern Instruments, Malvern, United Kingdom) instrument equipped with a 4 mW HeNe laser (633 nm) and a light scattering detector positioned at 90 °.

#### Bioactivity test in simulated body fluid

An *in vitro* bioactivity test was performed to evaluate the apatite-forming ability of SD-MBG and SD-MBG + BMP-2 in simulated body fluid (SBF) and prepared as previously published [67] (all components: Sigma Aldrich, St. Louis, USA). According to the protocol described by Maçon et al. [67], 2 mg of SD-MBG/ + BMP-2 were soaked in 2 mL of SBF (final concentration 1 mg/mL). The samples were kept soaked at 37 °C for up to 7 days on an orbital shaker (Excella E24, Eppendorf, Hamburg, Germany) with an agitation rate of 150 rpm. At each pre-defined time point (1 day, 3 days and 7 days), the suspension was centrifuged at 10,000 rpm for 5 min, in order to collect the powder. The pH of each recovered supernatant was measured, and the powder was washed with distilled water and dried in oven at 70° C overnight prior FE-SEM observations and EDS analysis to evaluate the apatite layer formation.

#### *In vitro* BMP-2 release experiments

*In vitro* release experiments were carried out in separate vials (n>3), each containing 75 µg of SD-MBG + BMP-2 (66.7 µg BMP-2 / mg SD-MBGs). Elution medium was PBS (Gibco/ Thermo Fisher Scientific, Waltham, USA) or Tris-HCl (Trizma Base, Sigma-Aldrich, St. Louis, USA), pH was set to 7.4, and the release experiment was carried out in 1 mL volume per vial at 37 °C. At each testing time point, the vials were centrifuged at 1000 g for 5 min, 500 µL elution medium was collected and stored at −20 °C, the SD-MBG vials were re-supplemented with fresh elution buffer. Quantification of released BMP-2 was accomplished using a α-BMP-2 enzyme-linked immunosorbent (ELISA) assay (Peprotech, Hamburg, Germany) according to the manufacturer’s instructions, testing each sample at least in duplicate. Using TMB as substrate, the optical density was measured at 450 nm with a reference measurement at 620 nm and the cumulative release was calculated. To adjust the amount of cumulative BMP-2 release to the dosage applied *in vivo*, the obtained values were multiplied by the factor 10 (0.75 mg of loaded material were applied in the fracture gap). BMP-2 has a described solubility in aqueous solutions of 0.1 – 1 mg/mL (Peprotech, Hamburg, Germany), therefore allowing this multiplication.

### *In vitro* cell culture experiments

#### Isolation and culture conditions

Primary human bone marrow mesenchymal stromal cells (hMSCs) from three different donors (female, aged 48 to 58, without known comorbidities) were included in the *in vitro* experiments. Human MSCs were obtained from the Core Facility “Tissue Harvesting of the BIH Center for Regenerative Therapies (BCRT)”. Metaphyseal bone marrow biopsies of patients routinely undergoing hip replacement surgeries at the Charité - Universitätsmedizin Berlin, Germany, were used for the isolation of hMSCs, which were afterwards characterized as described previously [68]. Tissue sampling and cell isolation were approved by the local Ethics Committee/ Institutional Review Board (IRB) of the Charité Universitätsmedizin Berlin, Germany, and written consent of the donor was obtained. Cells were cultured at 37 °C in 5 % CO_2_ atmosphere in Dulbecco’s modified Eagle’s medium (DMEM, low glucose) (Sigma Aldrich, St. Louis, USA) supplemented with 10 % v/v fetal bovine serum (FBS) (FBS Superior, Biochrom, Berlin, Germany), 1 % v/v GlutaMAX (Thermo Fischer Scientific, Waltham, USA), and 1 % v/v penicillin/streptomycin (Biochrom, Berlin, Germany), in the following termed expansion medium (EM). At a confluency of 70-80 %, cells were sub-cultured using TrypLE (Thermo Fischer Scientific, Waltham, USA) for cell detachment, cells up to passage 4 were used for all experiments. The experiments were carried out in 24-well tissue culture treated plates. For cell viability and cell number experiments, cells were seeded at a density of 2400 cells/cm^2^, for differentiation assays at a density of 6400 cells/cm^2^ in EM. After overnight attachment, the assays were started by exposing the hMSCs to ionic dissolution products from SD-MBG via adding the MBG to a transwell insert (6.5 mm Transwell^®^ with 0.4 µm pore size, Corning, Corning, USA), thereby exposing the cells only to the continuously generated ionic dissolution products of SD-MBG while avoiding direct physical interaction of hMSCs and SD-MBG. Two SD-MBG concentrations (1.5 and 5 mg/mL) were tested. Medium for cell expansion and osteoinductive experiments was changed two times per week, final volume per well was 500 µL. If not stated otherwise, assays were carried out according to the manufacturers’ instructions. Absorbance or fluorescence measurements were conducted using the Infinite 200 Pro and the Tecan i-control software (both: Tecan Group, Männedorf, Switzerland).

#### Metabolic activity, cell number and cytotoxicity under expansion conditions

The metabolic activity of hMSCs in EM was determined at day 1, 3 and 5 using Presto Blue (1:10 dilution in EM, 1h of incubation at 37°C, ex/em 560/590, Thermo Fisher Scientific, Waltham, MA). After 10 min of fixation using 4% neutral buffered formaldeyde (VWR, Darmstadt, Germany), the cell count was assessed by DAPI-staining (Sigma Aldrich, St. Louis, USA) of the nuclei (1 µg/mL, 15 min of incubation, washing with PBS, ex/em 358/ 461). Moreover, two 3×3 mosaic fluorescence images of the DAPI-stained wells were taken per well, cell number was counted using the software Fiji ImageJ [69]. To investigate cytotoxicity, the supernatants at day 1, 3 and 5 were collected, centrifuged and 25 µL of cell-/ debris-free supernatant was used per well for the LDH assay (Roche, Basel, Switzerland).

#### Osteogenic differentiation, phosphate and mineralization assays

To induce osteogenic differentiation, hMSCs were cultivated in 500 µL osteogenic differentiation medium (OM), which consists of expansion medium supplemented with 100 nM Dexamethasone, 50 µM L-Ascorbic acid 2-phosphate sesquimagnesium salt hydrate, and 10 mM ß-Glycerophosphate disodium salt hydrate (all: Sigma Aldrich, St. Louis, USA). Prior to each medium change, supernatants were collected, centrifuged and cell-/debris-free supernatants were stored at −80°C. The OM (+/-exposure to SD-MBG ionic dissolution products) supernatants were diluted 1:700, the EM controls 1:100 to be used in the phosphate assay (Abcam, Cambridge, United Kingdom). At 10 and 14 days of cultivation in OM, cells were washed with PBS, fixed in 4% neutral buffered formaldehyde (VWR, Darmstadt, Germany), cell number was determined by DAPI staining (Sigma-Aldrich, St. Louis, USA) as mentioned above. 0.5% w/v Alizarin Red S (Sigma-Aldrich, St. Louis, USA, 10 min incubation, RT) in distilled water was used to stain the mineralized extracellular matrix (ECM), thorough washing with distilled water removed unbound Alizarin Red S. For quantification, 10% w/v cetylpyridinium chloride (Sigma-Aldrich, St. Louis, USA) in distilled water was added to the air-dried wells and kept on an orbital shaker for 45 min, followed by an absorbance measurement of the dissolved stain at 562 nm. As an additional method to visualize deposited hydroxyapatite, an OsteoImage Assay (Lonza, Basel, Switzerland) was carried out. After washing, a DAPI staining to allow detection of cell nuclei was performed as described above. The cell cultures were imaged using a fluorescent microscope (BZ-X810, Keyence, Osaka, Japan) at 10x magnification.

### *In vivo* bone healing study

#### Housing conditions, osteotomy surgery and study design

The bone regeneration potential of the different treatments was studied in a rat osteotomy model of delayed healing as previously described [34]. A total of 18 adult female Sprague-Dawley rats (aged >7 months, ≥300 g, Janvier Labs, Le Genest-Saint-Isle, France), that had more than three litters (ex-breeders) were included in this study. Rats were kept in small groups under obligatory hygiene standards and conventional housing conditions with controlled temperature set to 20 ± 2 °C, a light/dark period of 12 h and food and water being available *ad libitum*. All animal experiments were approved by the local animal protection authorities (Landesamt für Gesundheit und Soziales Berlin, Germany: G0258/18) and performed in accordance with the German Animal Welfare Act, the National Institutes of Health Guide for Care and Use of Laboratory Animals and the ARRIVE guidelines.

Before starting the surgery, the rats were anesthetized by inhalation of isoflurane (Forene, Abott, Wiesbaden, Germany) and received a potent analgesic (Bubrenorphine, RB Pharmaceuticals, Berkshire, United Kingdom; 0.1 mg/kg BW), an antibiotic bolus (Clindamycin, Ratiopharm, Ulm, Germany; 45 mg/kg BW) and eye ointment. The osteotomy was carried out under deep anesthesia on a heating plate set to 37°C. The operation area of the left femur was clipped and disinfected, the femur was exposed by a longitudinal skin incision and blunt preparation of the muscles. An external fixator (RatExFix, RISytem, Davos, Switzerland) was mounted on the femur, followed by creation of a 2 mm osteotomy using an oscillating saw (W&H, Bürmoos, Austria) and a saw guide. The wound was closed with sutures and the rats were returned to their cages. As post-operative analgesia, Tramadolhydrochloride (Grünenthal, Aachen, Germany; 0.5 mg/mL) was added to the drinking water for three days post-surgery.

At 2 and 4 weeks post surgery, the animals were radiologically examined by X-rays and *in vivo* µCT (for details, see section below) under anesthesia induced by intraperitoneal injection (i.p.) of ketamine hydrochloride (Actavis Switzerland, Regensdorf, Switzerland; 60 mg/kg BW) and medetomidine (CP-Pharma, Burgdorf, Germany; 0.3 mg/kg BW). To recover from anesthesia, an antidote (1.5 mg/kg BW atipamezol, CP-Pharma, Burgdorf, Germany) was injected intra-muscularly (i.m.) for the 2 weeks time point. The final study time point was set to 4 weeks post surgery. After *in vivo* radiological examinations, blood was collected by intracardiac puncture, the animals were euthanized under deep anesthesia by intracardiac injection of potassium chloride, and the osteotomized femur was harvested. The bones were fixed in 4 % PFA (Science Services, München, Germany) in PBS for 24 h at 4 °C. The bones were again imaged using a higher resolution µCT, and afterwards dehydrated and paraffin-embedded for histological analysis.

#### Micro-computed x-ray tomography (µCT) analysis

At 2 and 4 weeks post osteotomy, the animals were radiologically examined by X-rays and *in vivo* µCT (Viva 40, SCANCO Medical, Wangen-Brüttisellen, Switzerland) under anesthesia. µCT nominal resolution was set at 35 µm voxel soze, with 55 kV source voltage and 145 µA source current. A global threshold was applied to all bones corresponding to a bone mineral density of 408 mg/cm^3^ calcium hydroxyapatite (CaHA). Four weeks post osteotomy, bones were harvested and cleaned of excess of soft tissue, fixed in 4 % PFA/PBS for 24 h and rinsed thoroughly in PBS. Harvested bones were mechanically fixed within a radiologically transparent serological pipette to keep the integrity of the bone and immersed in PBS. µCT scans were then performed using a Bruker SkyScan 1172 high-resolution micro-CT (Bruker, Kontich, Belgium) with a nominal resolution of 8 µm, 0.5 mm aluminum filter, 80 kV source voltage and 124 µA source current. A camera pixel binning of 2 × 2 was used together with an orbital scan of 180 degrees in steps of 0.3 degrees. Reconstruction was performed using the SkyScan NRecon software. Gaussian smoothing, ring artifact reduction, misalignment compensation and beam hardening correction were applied. The volume of interest (VOI) was defined to include the 2 mm defect region and 1 mm in the proximal and distal direction from the cutting plane of the bone defect. A global threshold was applied to all bones corresponding to a bone mineral density of 435 mg/cm^3^ CaHA. Calibration was performed using phantoms containing 0.25 and 0.75 g/cm^3^ CaHA (Bruker, Kontich, Belgium) homogenously distributed in epoxy rods of similar diameter as of the scanned bones to minimize beam hardening error.

#### Histological and immunohistochemical analysis

Paraffin-embedded bones were sectioned into 5 µm-thick sections, de-paraffinized by 2x 10 min of incubation in Xylol and re-hydrated by descending alcohol series and distilled water as the final step before staining, stained for MOVAT’s Pentachrome as previously described [70] and for Sirius Red (Sigma-Aldrich, St. Louis, USA). Briefly, slides were incubated for 1 h in 1 % Sirius Red solution and washed twice with 0.5 % acetic acid (Sigma-Aldrich, St. Louis, USA). Afterwards, slides were washed in 1 % acetic acid. At the end of the staining, the slides were dehydrated using Xylol (Fisher Chemical, Thermo Fisher Scientific, Waltham, USA) and embedded with Vitroclud (Langenbrink, Emmendingen, Germany). For the immunohistochemical analysis, after deparaffinization and re-hydration, slides were blocked using 5 % normal horse serum (Vector Laboratories, Burlingame, USA) for 1 h and 1 % BSA/PBS, followed by overnight incubation at 4 °C with α-α-SMA (1:400, mouse monoclonal, clone 1 A4, DAKO Agilent Technologies, Santa Clara, USA) or α-CD68 (1:2000, mouse monoclonal, clone BM4000, OriGene Technologies, Rockville, USA). An α-mouse, rat adsorbed biotinylated secondary antibody (Vector Laboratories, Burlingame, USA) diluted 1:50 in 2 % normal serum horse and 1 % BSA/PBS was incubated on the slides for 30 min. AB complex (Vector AK 5000, Vector Laboratories, Burlingame, USA) was incubated for 50 min, then the milieu was slightly alkalized by using a chromogen buffer (pH 8.2), followed by the visualization of the staining (Vector SK 5100, Vector Laboratories, Burlingame, USA). As counterstaining, hematoxylin (Mayer’s) was chosen and the slides were embedded using Aquatex (Merck, Darmstadt, Germany). Microscopic images of all slides were taken at 10x magnification under bright field (Axioskop 40, Carl Zeiss, Oberkochen, Germany). Histomorphometric analyses were carried out using the MOVAT’s pentachrome-stained slides and applying a custom-made macro embedded in FIJI ImageJ Software [69]. Callus area was determined manually. Detection of mineralized tissue and cartilage was performed according to color thresholding to determine the areas of the respective tissues. Blood vessels and osteoclasts were revealed in an analog manner by α-SMA and CD68 staining respectively. Finally, the amount of blood vessels was normalized to the total area of the callus and the length of mineralized callus surface covered by CD68+ cells to was normalized to the total length/ perimeter of the mineralized callus.

### Statistics

The statistical evaluation of the presented data was performed using GraphPad Prism® (GraphPad Software, San Diego, USA). Confidence interval was set to 0.95, p-values for statistical significance were \**p* < 0.05, \*\**p* < 0.01, \*\*\**p* < 0.001, ****p < 0.0001. Detailed information on all statistical analyses performed, including statistical tests, depicted values and sample size, are mentioned in the figure legends. In general, for the small sample sizes of the *in vivo* preclinical study, the data can not be considered normally distributed. Accordingly, the statistical test applied was a Mann-Whitney U test. For larger sample sizes that were tested for normal distribution, an ANOVA with Tukey’s multiple comparison test was performed.

## Acknowledgement

We thank Sabine Stumpp and Norma Schulz (Julius Wolff Institut, Charité - Universitätsmedizin Berlin) for assistance in the histological preparation of the rat bone sections and Gabriela Korus (Julius Wolff Institut, Charité Universitätsmedizin Berlin) for the support in setting up the immunohistological staining protocol. We moreover wish to express our gratitude to the Core Facility “Tissue Harvesting of the BIH Center for Regenerative Therapies (BCRT)” for providing the hMSCs used in the *in vitro* studies. This project has received funding from the European Union’s Horizon 2020 research and innovation program under grant agreement No. 685872-MOZART (www.mozartproject.eu).

