## Supplements for "*In vivo* validation of spray-dried mesoporous bioactive glass microspheres acting as prolonged local release systems for BMP-2 to induce bone regeneration"

#### Absorbable collagen sponge impairs healing demonstrated by *ex vivo* µCT images

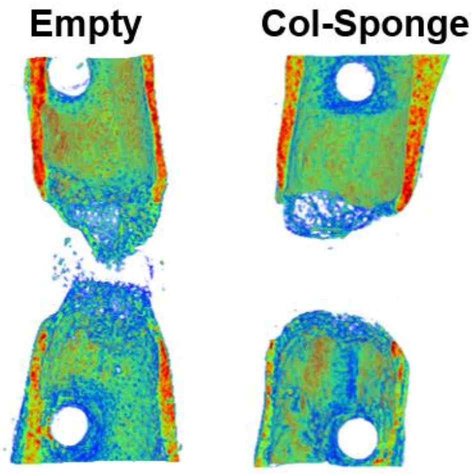


**Figure S1. Commercially available collagen sponge impairs bone healing.** Reconstructed representative *ex vivo* µCT images of osteotomized murine femora (1.4 mm fracture gap) 21 days post osteotomy comparing the bone formation of animals without addition of any material into the gap (empty) and animals that received a collagen sponge (Lyostypt, bovine collagen-1, B. Braun, Germany). The formation of new bone is stronger in the empty than in the collagen sponge group.

#### Human whole blood assay and quantification of immune reaction via human α-TNF-α/ α-IFN-γ ELISA

<u>Method</u>

For the human whole blood assay, heparinized human peripheral blood was withdrawn from a healthy donor. SD-MBG were dispersed in RPMI1640 (Biochrom, Berlin, Germany) without additives and mixed 1:2 with whole blood in a 24-well culture plates, allowing direct contact of material and immune cells. The tested SD-MBG concentration was 5 mg/ml. For comparison, whole blood was stimulated with Lipopolysaccharide (LPS, O111:B4, Sigma Aldrich, St. Louis, USA) at concentrations of 0.1, 10 and 500 ng/ml. At 4 and 24 h post addition of the SD-MBG or LPS, respectively, the plates were centrifuged to generate debris-free supernatant, the supernatant was harvested and stored at -80°C until ELISA-based cytokine quantification. Human α-TNF-α and α-IFN-γ (Invitrogen/ Thermo Fisher Scientific, Waltham, USA) were carried out according to the manufacturer's instructions; each sample was tested in duplicate.

<u>Results</u>

At 4 h post addition of the SD-MBG or LPS to human whole blood, the immune reaction against the material can be considered weak with TNF-α levels of around 35 pg/ml, as seen by the almost 200 times higher secretion of this cytokine under low-level stimulation with 0.1 ng/ml LPS. At 24 h, the TNF-α secretion due to SD-MBG diminished entirely. IFN-γ was found not secreted in response to the SD-MBGs at both time points.

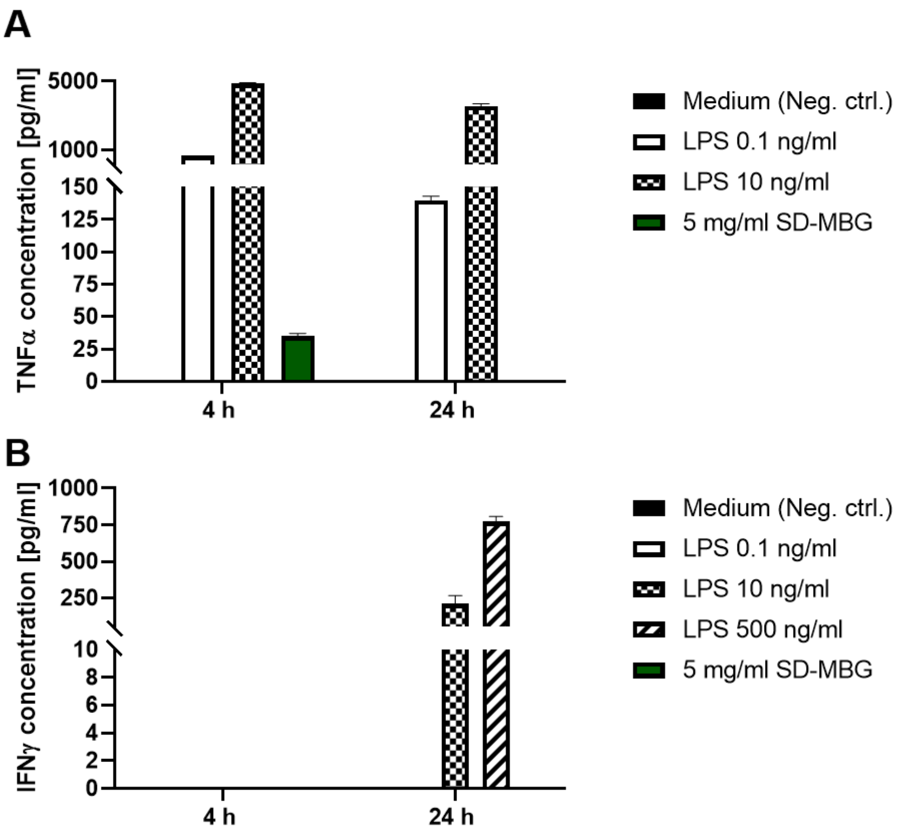


**Figure S2. Quantification of pro-inflammatory cytokine secretion by immune cells contained in human whole blood after exposure to SD-MBG for 4 and 24 h. A:** TNF-α and **B:** IFN-γ secretion, quantification shown for three replicates of material exposure to whole blood of one donor. For comparison, LPS was used in different concentrations. N= 2 for the LPS control groups, n= 3 for SD-MBGs supplemented group. All samples measured in duplicate in the ELISA.

#### Osteogenic differentiation capacity testing – comparison of 5 donors

HMSCs from five different donors were tested for their *in vitro* osteogenic differentiation capacity by inducing osteogenesis and performing an Alizarin Red S staining and cell nuclei staining (DAPI) at 10 and 14 days post osteogenic induction. Please see Materials and Methods section for experimental details. Based on the amount of deposited extracellular hydroxyapatite, hMSCs from three donors showing limited osteogenic capacity were selected for subsequent experiments with SD-MBG, since we aimed at unraveling beneficial effects of the SD-MBGs application which could be concealed in highly mineralizing hMSCs.

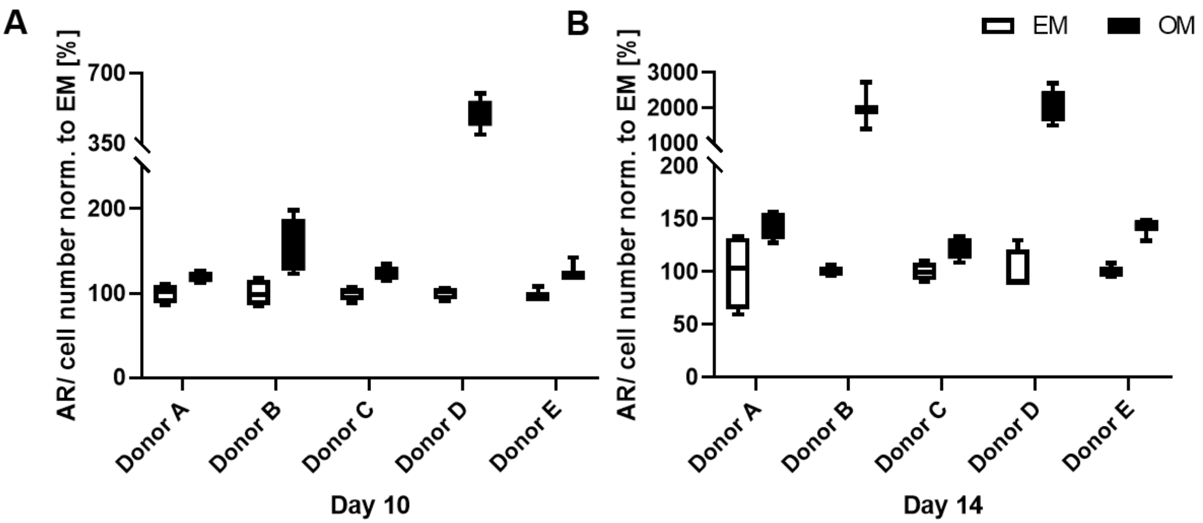


**Figure S3. Testing of osteogenic differentiation capacity on human primary hMSCs.** Increase in mineralization potential due to osteogenic induction (OM) was determined by measuring the optical density of Alizarin Red S Staining (AR) per cell number determined by DAPI staining and normalized to the control cells kept in expansion medium (EM) at 10 (**A**) and 14 (**B**) days. Note that donor B and D show substantial matrix mineralization indicating a strong intrinsic osteogenic differentiation potential of these hMSCs. N= 5 different donors of hMSCs tested in ≥3 technical replicates.

### SD-MBG material characterization by nitrogen adsorption analysis

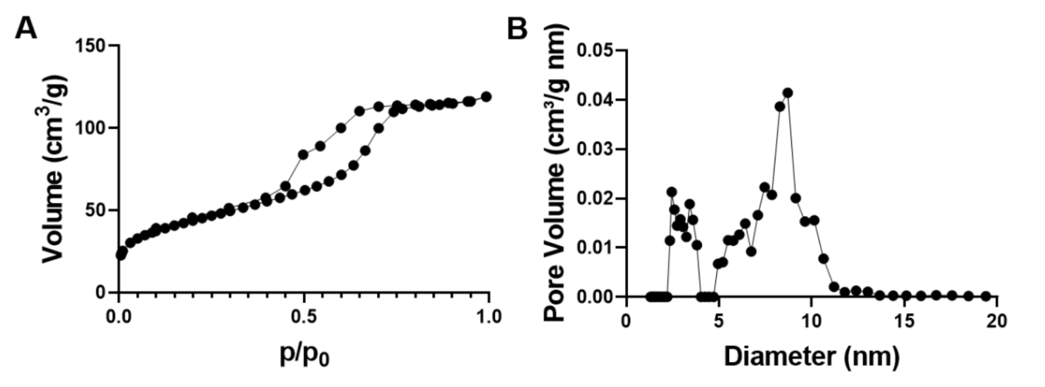


**Figure S4. N2 adsorption-desorption isotherm (A) and DFT pore size distribution (B) of SD-MBG.** $N_2$ adsorption-desorption measurement was conducted by using an ASAP2020, Micromeritics analyser at a temperature of –196 °C; before the analysis, samples were outgassed at 150 °C for 5 h. The Brunauer-Emmett-Teller (BET) equation was used to calculate the specific surface area ($SSA_{BET}$) from the adsorption isotherm in the 0.04–0.2 relative pressure range. The pore size distribution was calculated by using the DFT method (Density Functional Theory). $N_2$ adsorption-desorption isotherm of SD-MBGs is an IV type isotherm with a pronounced hysteresis loop, characteristic of mesoporous materials.

### *In vivo* µCT

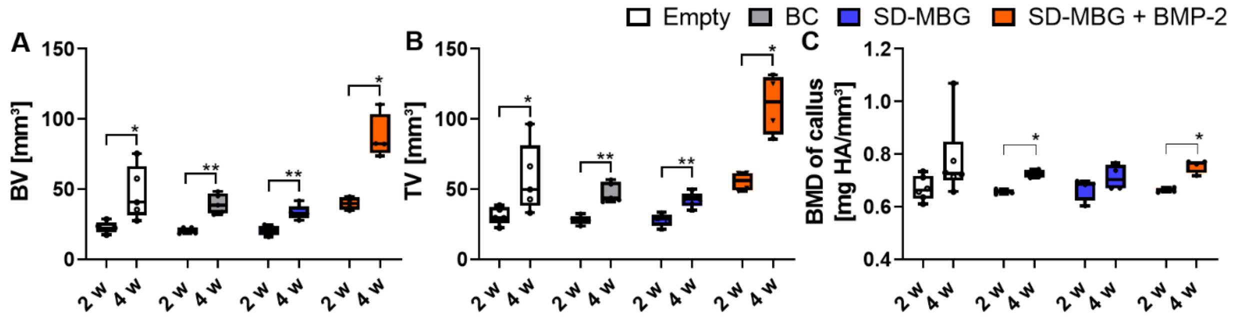


**Figure S5. *In vivo* µCT analysis. A:** bone volume (BV) and **B:** tissue volume (TV) at 2 and 4 weeks post osteotomy. To compare the increase in BV and TV with progressing healing time, a Mann-Whitney-U test was performed per group, always comparing the 2 with the 4 weeks time point. **C:** Bone mineral density (BMD) of the newly formed callus, excluding the cortical bone fragments that are part of the region of interest at 2 and 4 weeks post surgery. For the comparison of the development of BMD over time per group, a Mann-Whitney-U test was performed separately for each group. To understand if differences in the BMD between

BMP-2 and other groups at each time point occurred, a Mann Whitney U test was performed using the BMP-2 group as comparator, yielding no statistically significant differences. N= 4-6 animals per group, shown are box plots min and max with a line at the median.

#### Examples for reconstructed µCT images in two planes for the evaluation of bridging

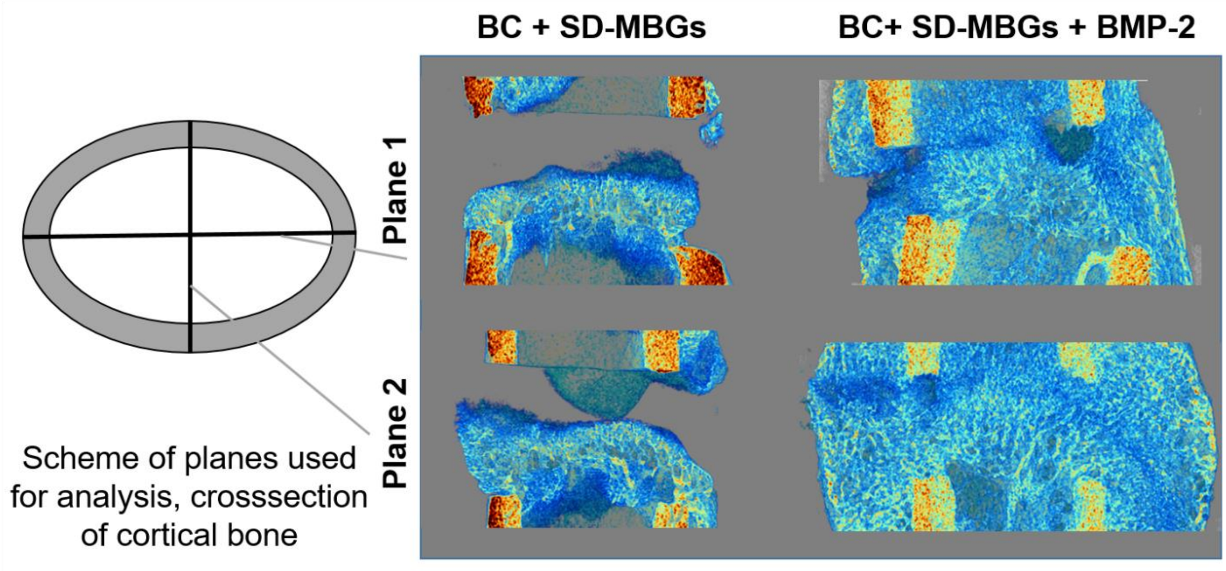


**Figure S6. Evaluation of bridging based on reconstructed µCT images cut longitudinally in two planes.** Schematic representation of the two planes used for the evaluation. Representative images of the BC + SD-MBG with (right) and without (left) BMP-2; the same bone is shown cut along both planes as indicated in the schematic figure.

### Phosphorous and Calcium abundance in the fracture hematoma of sheep

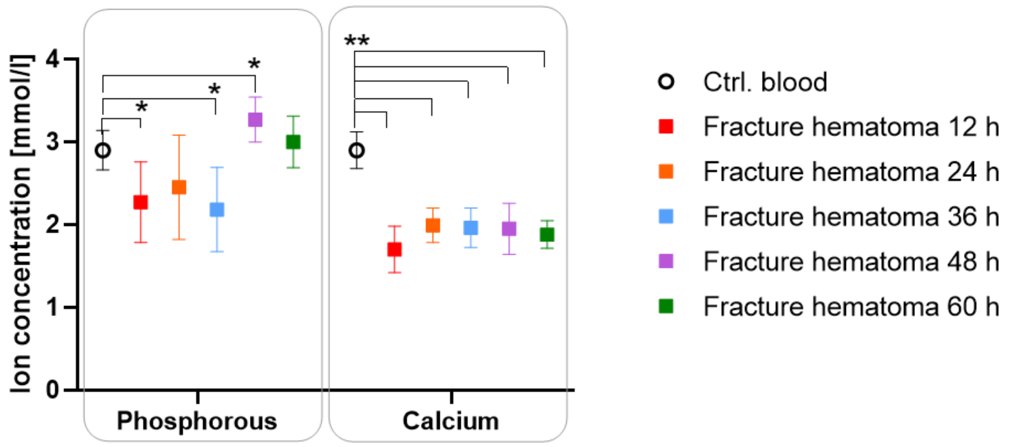


**Figure S7. Determination of ion species abundance for Phosphorous and Calcium in sheep fracture hematomas at 12, 24, 36, 48 and 60 hours post osteotomy.** For comparison, peripheral blood was collected prior to surgery. N=6 per time point. Per ion, a Mann-Whitney U test was performed, using the control blood as comparator.
